# Cellulonodin-2 and Lihuanodin: Lasso Peptides with an Aspartimide Post-translational Modification

**DOI:** 10.1101/2021.05.19.444711

**Authors:** Li Cao, Moshe Beiser, Joseph D. Koos, Margarita Orlova, Hader E. Elashal, Hendrik V. Schröder, A. James Link

## Abstract

Lasso peptides are a family of ribosomally synthesized and post-translationally modified peptides (RiPPs) defined by their threaded structure. Besides the class-defining isopeptide bond, other post-translational modifications (PTMs) that further tailor lasso peptides have been previously reported. Using genome mining tools, we identified a subset of lasso peptide biosynthetic gene clusters (BGCs) that are colocalized with protein L-isoaspartyl methyltransferase (PIMT) homologs. PIMTs have an important role in protein repair, restoring isoaspartate residues formed from asparagine deamidation to aspartate. Here we report a new function for PIMT enzymes in the post-translational modification of lasso peptides. The PIMTs associated with lasso peptide BGCs first methylate an L-aspartate sidechain found within the ring of the lasso peptide. The methyl ester is then converted into a stable aspartimide moiety, endowing the lasso peptide ring with rigidity relative to its unmodified counterpart. We describe the heterologous expression and structural characterization of two examples of aspartimide-modified lasso peptides from thermophilic Gram-positive bacteria. The lasso peptide cellulonodin-2 is encoded in the genome of actinobacterium *Thermobifida cellulosilytica*, while lihuanodin is encoded in the genome of firmicute *Lihuaxuella thermophila*. Additional genome mining revealed PIMT-containing lasso peptide BGCs in 48 organisms. In addition to heterologous expression, we have reconstituted PIMT-mediated aspartimide formation *in vitro*, showing that lasso peptide-associated PIMTs transfer methyl groups very rapidly as compared to canonical PIMTs. Furthermore, in stark contrast to other characterized lasso peptide PTMs, the methyltransferase functions only on lassoed substrates.

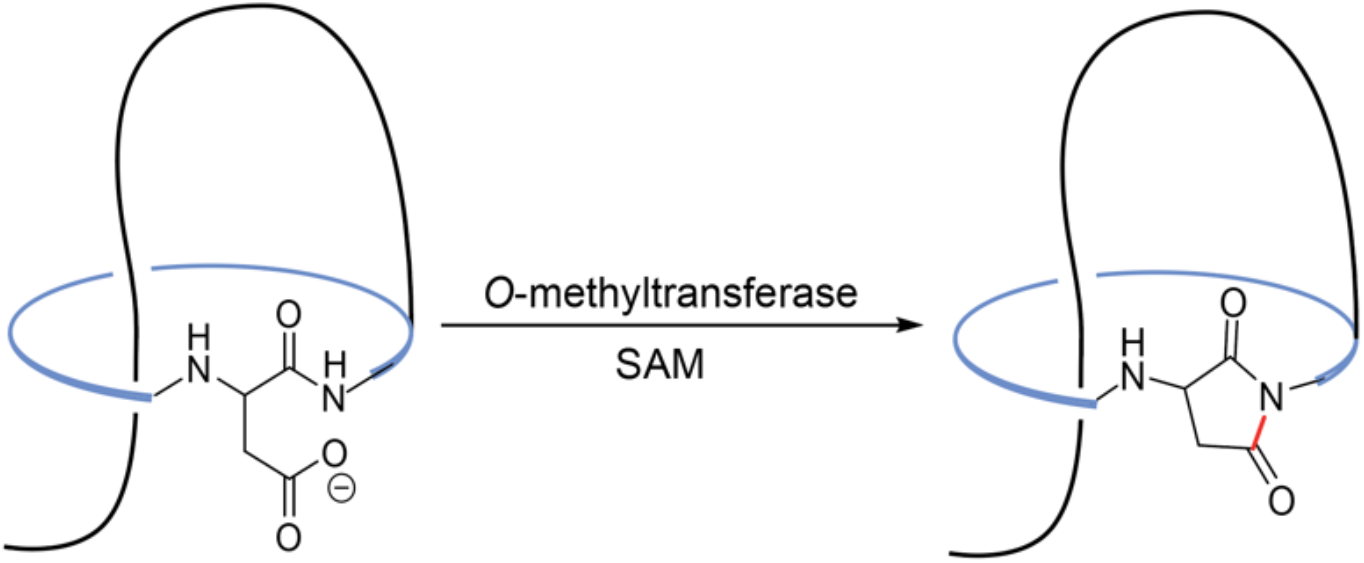

---

Lasso peptides are a family of ribosomally synthesized and post-translationally modified (RiPP) natural products.^1,2^ Structurally, lasso peptides are [1]rotaxanes, in which an isopeptide bond joins the N-terminus of the core peptide to the side chain of a D or E residue, forming a 7-9 amino acids (aa) macrocyclic ring.^3–5^ The C-terminal tail portion of the peptide, ranging from 6 to 25 aa,^6,7^ is threaded through the ring, resulting in a right-handed, lassoed three-dimensional conformation. The size of the lasso peptide loop, which extends from the isopeptide bond and threads through the ring, varies in size from as few as 4 aa up to 18 aa.^6,8^ Likewise, the C-terminal tail of the peptide can vary from as few as 2 aa up to 18 aa.^7,9^ Lasso peptide biosynthesis follows the general biosynthetic steps of other RiPPs.^10–16^ The precursor peptide, encoded by the A gene, is first processed by a cysteine peptidase, encoded by the B gene, which hydrolyzes the peptide bond between the leader and the core peptide. In many Gram-positive bacteria, the B gene is split between two open reading frames, named B1 and B2.^17^ The lasso is then formed via the action of the protein encoded by the C gene, an asparagine synthetase homolog, which is hypothesized to activate the Asp or Glu residue with ATP and install an isopeptide bond to template the lasso structure. Gene clusters encoding lasso peptides can also include additional genes besides the ones required for lasso peptide biosynthesis. Of the characterized lasso peptide biosynthetic gene clusters (BGCs), examples that encode antimicrobial lasso peptides also contain a D gene that encodes an ABC transporter, which is dedicated to exporting the matured peptide and serves as an immunity factor.^11,18–23^ Another group of BGCs contains a conserved E gene encoding an isopeptidase that specifically cleaves the isopeptide bond of lasso peptides encoded by the neighboring genes.^3,17,24,25^

Beyond the minimal BGC required for lasso peptide biosynthesis and commonly associated transporters and isopeptidases, genome mining of the ever-growing database of bacterial genome sequences has revealed tailoring enzymes that subject lasso peptides to further post-translational modifications (PTMs). Modifications that have been confirmed experimentally include disulfide bonds,^26,27^ C-terminal methylation,^19,28^ acetylation,^29^ citrullination,^30^ phosphorylation,^31^ glycosylation,^32^ epimerization,^26,33^ and *β*-hydroxylation.^6,34^ All previously described lasso peptide PTMs occur in either the loop or tail regions of the lasso peptide, and these auxiliary enzymes recognize the linear precursor rather than the matured lasso as a substrate. ^31,34^

Here we report the discovery of two lasso peptides, cellulonodin-2 from actinobacterium *Thermobifida cellulosilytica*, and lihuanodin from firmicute *Lihuaxuella thermophila*. The BGCs for cellulonodin-2 and lihuanodin include a gene annotated as an *O*-methyltransferase which has homology to protein L-isoaspartyl methyltransferases (PIMTs). The canonical role of PIMTs is reversion of isoaspartyl (isoAsp) residues that result from protein aging back to L-aspartate.^35–37^ Methylation of isoAsp by the PIMT leads to non-enzymatic formation of a succinimide moiety, aspartimide, which is then hydrolyzed to a mixture of isoAsp and Asp. Repeated catalytic cycles result in conversion of most of the isoAsp residues to Asp. In contrast to canonical PIMTs, the PIMT homologs in the cellulonodin-2 and lihuanodin BGCs act on Asp. Compared to characterized PIMTs, the *O*-methyltransferases associated with these lasso peptide BGCs are exceptionally fast examples of methyl esterification catalysts. Furthermore, the aspartimide residue appears to be the intended final product, rather than just an intermediate. The modification of cellulonodin-2 and lihuanodin occurs at an Asp residue within the lasso peptide ring, part of a strongly conserved DTAD motif. Thus, these peptides are the first examples of lasso peptides with PTMs in the ring. We also show here that the *O*-methyltransferases only recognize matured lassoes, not linear precursors, as their substrates. This is in contrast to other lasso peptide PTMs which are carried out on the precursor.

## Results

### Genome Mining Reveals Lasso Peptide BGCs with *O*-Methyltransferase Genes

Our investigation into *Thermobifida cellulosilytica* began when two lasso peptide gene clusters were identified using our precursor-centric genome mining algorithm.^38^ The first cluster encoded the lasso peptide cellulonodin-1, whose core sequence and processing enzymes were highly similar in sequence to those required for the biosynthesis of lasso peptide fuscanodin (also known as fusilassin) from phylogenetically related *Thermobifida fusca* (8/18 identities in the core, Figure S1).^10,39^ The second cluster exhibited the *A-C-B1-B2* gene architecture that was typical of actinobacterial lasso peptide BGCs, with low sequence similarity as compared to the fuscanodin and cellulonodin-1 BGCs (Figure S1). In addition, the cluster also contained a gene (WP_068755506, 1023 bp) annotated as “methyltransferase domain-containing protein” (Figure 1a). A BLAST search on the putative translated sequence of this gene yielded many other methyltransferases, with some annotated as protein L-isoaspartyl *O*-methyltransferase (PIMT). Of these methyltransferases, 48 were found adjacent to putative lasso peptide BGCs, with 43 from actinobacteria and 5 from firmicutes. A sequence similarity network analysis on the *T. cellulosilytica* methyltransferase revealed that lasso peptide-associated methyltransferases were clustered together (Figure S2).^40^ Genes encoding methyltransferases appeared primarily as the last open reading frame (ORF) in the cluster, constituting the *A-C-B1-B2-M* gene architecture identical to the one in *Thermobifida cellulosilytica* (Figure 1a), where the *M* gene encodes the PIMT homolog. Exceptions existed in the BGCs from firmicutes, such as the one from *Lihuaxuella thermophila*, where *the M* gene was the first ORF instead, constituting an *M-A-C-B1-B2* architecture (Figure 1a). *O*-methyltransferases are also found in the BGCs for lassomycin-like lasso peptides,^28^ where the enzyme esterifies the C-terminus of these peptides. We carried out homology modeling of the PIMT homologs from *T. cellulosilytica* (TceM) and *L. thermophila* (LihM). These models showed essentially no structural similarity to the homology model for the methyltransferase in lassomycin-like BGCs (Figure S3).

**Figure 1:**
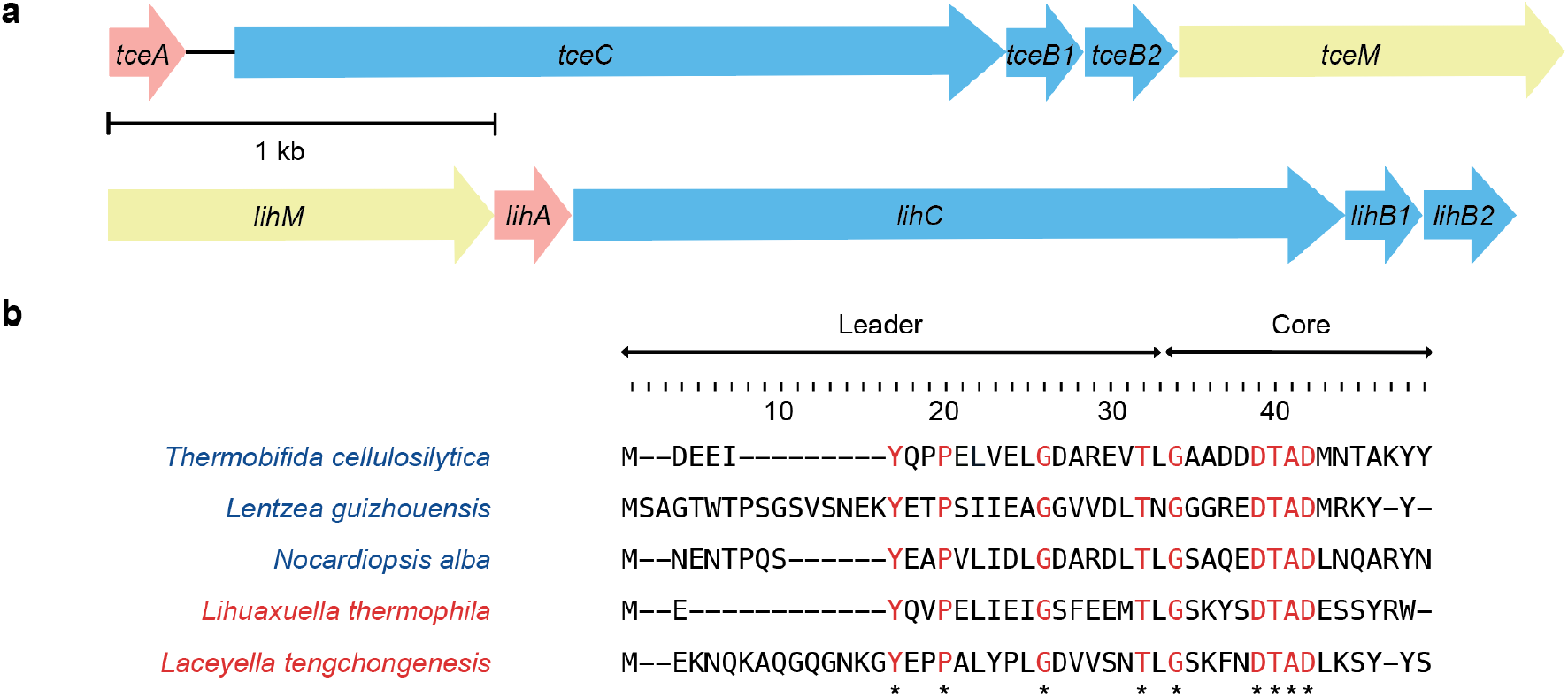
Lasso Peptide Gene Clusters Bearing a Protein L-Isoaspartyl Methyltransferase (PIMT) Homolog. **a)** Native lasso peptide biosynthetic gene clusters from actinobacterium *Thermobifida cellulosilytica* and firmicute *Lihuaxuella thermophila*. The *A, C, B1*, and *B2* genes are the minimal machinery for lasso peptide biosynthesis and encode the precursor peptide, the lasso cyclase, and the bipartite cysteine peptidase, respectively. The *M* genes, characterized here for the first time, are *O*-methyltransferases. The *tceM* gene is the last ORF in the cluster, while *lihM* is the first ORF. Clusters are drawn to scale; note the 126 base pair intergenic region between *tceA* and *tceC*. **b)** Sequence alignment of 5 selected putative lasso peptide precursors. Organisms labelled blue are actinobacteria, while those labelled red are firmicutes. The leader and core region of the precursor are indicated. Conserved residues are marked with asterisks and labeled red. In the core peptide region, there is a conserved G at position 1, and a conserved DTAD motif at position 6–9.

Aligning precursor sequences of 48 lasso peptide BGCs with PIMT homologs revealed a number of strongly conserved sequences, including the essential T residue in the penultimate position in the leader,^41^ conserved Y, P, and G residues in the leader region,^42^ a G residue at position 1, and a putative isopeptide bond forming D at position 9 (Figure 1b, Figure S4). Moreover, a highly conserved DTAD motif was observed in the predicted ring of putative lasso peptides (Figure 1b, Figure S4). Particularly, D6 is strictly conserved in all the putative lasso peptide precursors, suggesting it could be the site of modification by the PIMT homolog. Based on our previous success with heterologous expression of BGCs from thermophiles,^10^ we decided to pursue putative lasso peptide BGCs from two thermophiles, *Thermobifida cellulosilytica* and *Lihuaxuella thermophila*, as one example from actinobacteria, and the other from firmicutes.

### Heterologous Expression of Cellulonodin-2 and Lihuanodin

Several different iterations of gene cluster refactoring were carried out for both lasso peptide BGCs (detailed in Supporting Information). In the most optimized version for the BGC from *T. cellulosilytica*, the precursor gene *tceA* was placed under an IPTG-inducible promoter, and genes encoding the biosynthetic enzymes and the methyltransferase *tceCB1B2M* were placed under the constitutive promoter upstream of the microcin J25 biosynthetic enzymes, p_*mcjBCD*_ (Figure 2a). The native ORF that encoded *tceB2* had a GTG start codon and was changed to ATG. In the most optimized version for the *L. thermophila* BGC, we refactored the cluster into a co-expression system. The precursor gene *lihA* was placed under an IPTG inducible promoter, and genes encoding the biosynthetic enzymes *lihCB1B2* were placed under p_*mcjBCD*_. The start codon of *lihC* was changed from TTG to ATG. Since *lihM* is the first ORF in the biosynthetic gene cluster, it was placed under an IPTG-inducible T7 promoter on a separate plasmid (Figure 2a).

**Figure 2:**
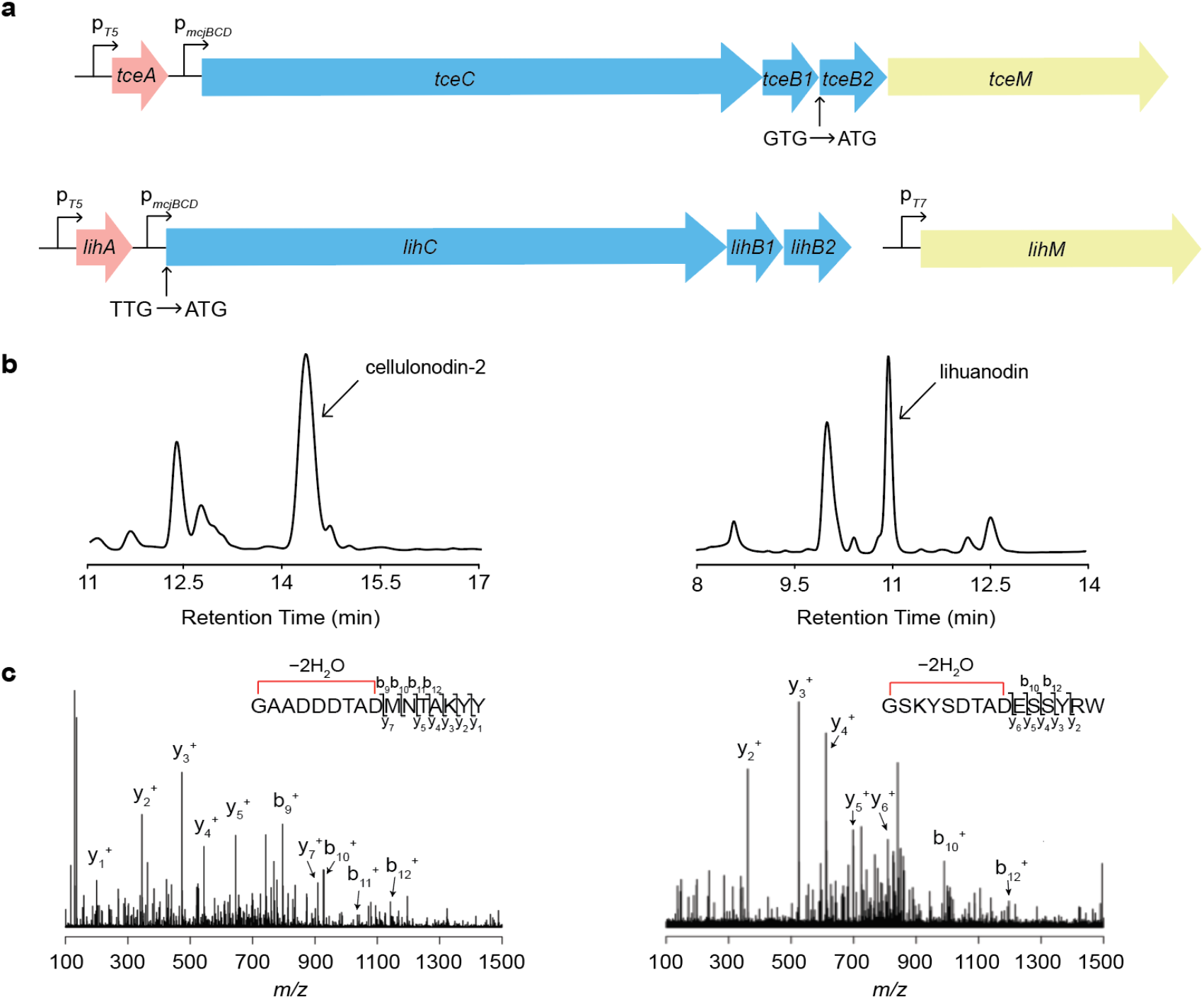
Heterologous Expression of Cellulonodin-2 and Lihuanodin Suggests an Additional PTM. **a)** Refactored gene clusters for cellulonodin-2 (top) and lihuanodin (bottom). IPTG-inducible T5 promoters were introduced upstream of both precursor peptide genes while the constitutive promoter from the MccJ25 BGC was used to drive expression of the maturation enzymes. In the lihuanodin expression system, an IPTG-inducible T7 promoter is used to drive methyltransferase (LihM) expression. All non-ATG start codons were reverted to ATG. **b)** HPLC trace of crude cell lysates containing cellulonodin-2 (left) and lihuanodin (right). **c)** Tandem mass spectra of cellulonodin-2 (left) and lihuanodin (right). The *y*-ions could be traced back all the way to the isopeptide-bonded ring. The N-terminal fragment contained two dehydrations, suggesting that an additional linkage was formed other than the isopeptide bond that defines lasso peptides.

Heterologous expression of both BGCs was carried out using *E. coli* BL21. The expected monoisotopic mass of the core peptide with isopeptide bond formation (i.e. a “lassoed only” species) was 1702.68 Da for the *T. cellulosilytica* lasso and 1732.74 Da for the *L. thermophila* lasso; the putative methylated species for these peptides have masses of 1716.69 Da and 1746.75 Da (Figure S5). In the cell extracts, there was a prominent peak that eluted at 14.1 min for the *T. cellulosilytica* BGC and 11.0 min for the *L. thermophila* BGC (Figure 2b). We collected these two peaks and found that they correspond to two species having monoisotopic masses of 1684.66 Da and 1714.72 Da, respectively. Both species correspond to an additional dehydration of the predicted isopeptide-bonded lasso peptides. We confirmed both species were indeed related to the predicted core peptide sequence by MS/MS (Figure 2c). The MS/MS results showed that there were two dehydrations within the first nine amino acids for both peptides, suggesting an additional bond formation other than the isopeptide bond that is a signature of lasso peptides. For the *T. cellulosilytica* lasso, lassoed only lassoed only and methylated products (1% and 14% by LC-MS peak area, respectively) were detected in the cell extract. For the *L. thermophila* lasso, small amounts of lassoed only and methylated species (2.8% and 1.7% by LC-MS peak area, respectively) were also detectable by LC-MS. Therefore, we named these putative lasso peptides with an additional dehydration cellulonodin-2 and lihuanodin, as cellulonodin-2 was the second lasso peptide discovered from *T. <u>cellulo</u>silytica*, and lihuanodin was named based on the genus in which it is encoded, *<u>Lihua</u>xuella*. Notably, expression of lasso peptides from firmicutes remains rare. Lihuanodin is only the third lasso peptide from firmicutes that has been experimentally verified.^31,32^ The yields of cellulonodin-2 and lihuanodin were 30 and 35 μg/L culture, respectively. Both peptides were only detected in *E. coli* cell pellets, and not in culture supernatants, consistent with the lack of an ABC transporter in these BGCs.

Additionally, constructs with only the minimal biosynthetic enzymes were assembled to confirm that TceM and LihM carried out the additional dehydration *in vivo*. These two constructs were also introduced into *E. coli* BL21, and the cell extracts were analyzed by HPLC and LC-MS. Only masses corresponding to lassoed only products without further PTMs were observed for both the cellulonodin-2 (1702.68 Da) and lihuanodin (1732.74 Da) BGCs. The identities of both species were also confirmed by MS/MS (Figure S6). These results suggested that the PTMs on both cellulonodin-2 and lihuanodin were not obligate for lasso formation.

Since both TceM and LihM were predicted to be PIMT homologs, we constructed *E. coli* BL21*Δpcm* to evaluate whether Pcm, the innate PIMT in *E. coli*, could alter the intended product *in vivo*. We heterologously expressed cellulonodin-2 in both *E. coli* BL21*Δpcm* and *E. coli* BL21, and LC-MS traces as well as product distributions remained the same in both cases (Figure S7), suggesting that the native PIMT in *E. coli* was not involved in the process. We used wild-type *E. coli* BL21 for all subsequent experiments.

### Cellulonodin-2 and Lihuanodin Contain an Unusually Stable Succinimide Moiety

We hypothesized that the additional dehydration in the ring segments of both cellulonodin-2 and lihuanodin was due to the presence of succinimide formed from an Asp residue, also known as aspartimide.^43^ As discussed above, aspartimide is an intermediate in protein repair pathways catalyzed by canonical PIMTs, but our data suggest that the aspartimide-modified lasso peptides are the final products of these BGCs. In this scenario, the PIMT homolog methylates an Asp sidechain, followed by nucleophilic attack of the adjacent amide, forming aspartimide (Figure 3a) The cofactor *S*-adenosyl methionine (SAM) serves as a methyl donor. An alternate reaction path that would lead to the observed dehydration in cellulonodin-2 and lihuanodin is attack on aspartyl methyl ester or aspartimide by Ser, Thr, or Lys sidechains, forming a new ester or amide linkage. To differentiate between these possibilities, hydrazine (2M, 35% in water) was added to 0.1 mM cellulonodin-2 in acetonitrile (ACN). A similar reaction was set up for lihuanodin, except that DMSO was used as a solvent due to poor solubility of lihuanodin in ACN. The reaction mixtures were incubated at room temperature for 30 min. If these peptides contained an aspartimide moiety, a mass change of +32 was expected (Figure 3b).^44^ In contrast, esters and amides were not expected to react with hydrazine under the test conditions.^45^ The starting materials reacted quantitatively under this condition, and peaks with a +32 mass change emerged for both peptides, suggesting both cellulonodin-2 and lihuanodin contained an aspartimide moiety (Figure 3c, d, Figure S8). Since we used a hydrazine solution in water, a small amount of hydrolyzed product was also observed in the case of lihuanodin (Figure 3d).

**Figure 3:**
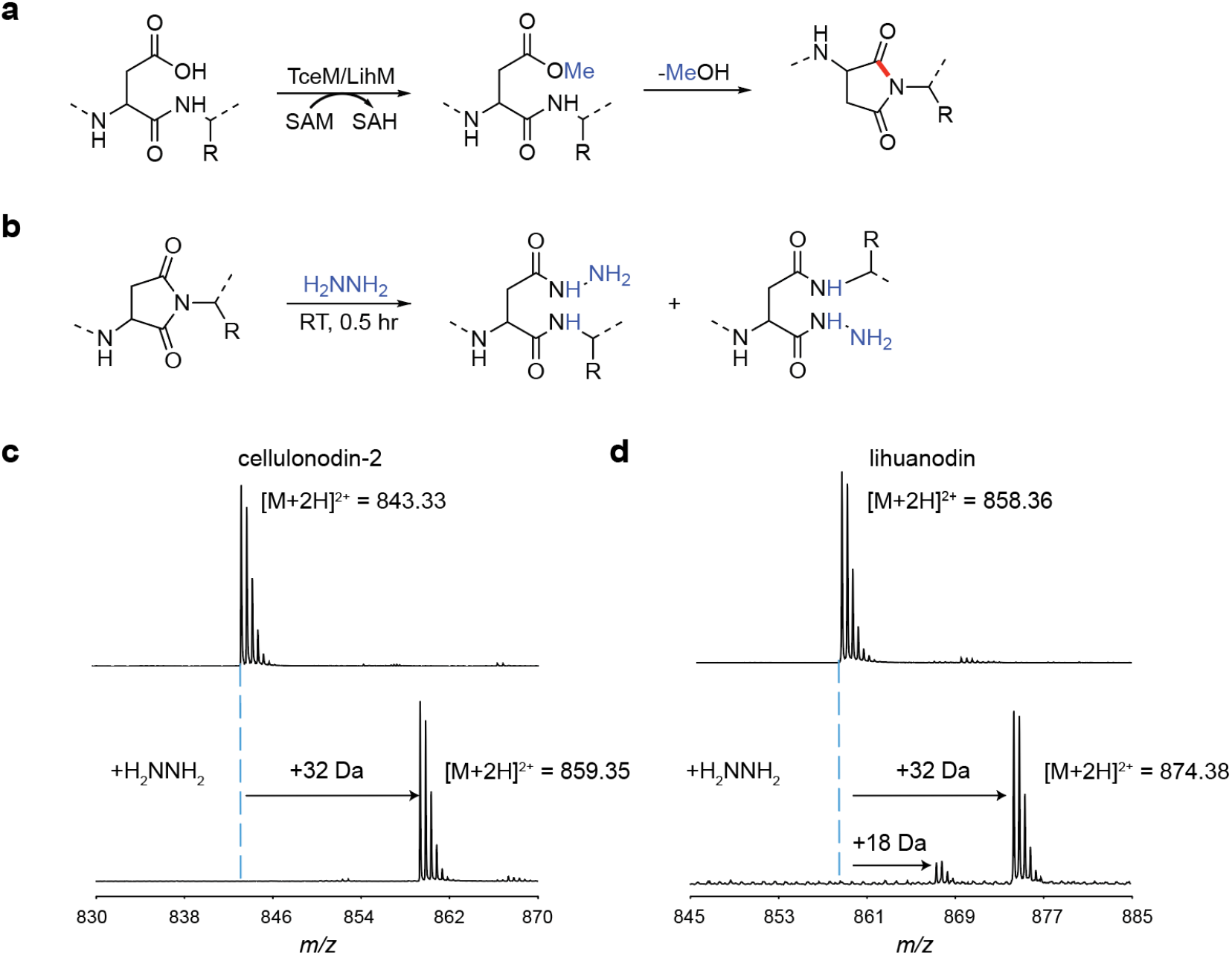
Cellulonodin-2 and Lihuanodin Contain Aspartimide. **a)** Proposed reaction pathway for TceM and LihM. The enzymes methylate aspartate (blue) followed by a putative non-enzymatic dehydration. The newly-formed bond is in red. **b)** Chemical scheme of a succinimide reacting with hydrazine. The hydrazine nucleophile can open the succinimide to give two different products. **c)** Mass spectrum of cellulonodin-2 reacted with hydrazine; the only products observed were the hydrazides. **d)** Mass spectrum of lihuanodin reacted with hydrazine. The hydrazides are the major product, with some minor hydrolysis.

Since both cellulonodin-2 and lihuanodin contained an aspartimide moiety, we were interested in determining the conditions under which this normally labile group could remain intact. We first incubated the HPLC-purified cellulonodin-2 and lihuanodin in ultrapure water at 20 °C, and analyzed the sample at 0, 2, and 22 h by LC-MS (Figure 4a). We observed no evidence of aspartimide hydrolysis under these conditions. For cellulonodin-2, a small amount of oxidized product was observed at 0 h (Figure 4a). The location of oxidation was confirmed to be on M10 by MS/MS (Figure S9). Upon incubation, a shoulder peak with the same mass as cellulonodin-2 grew over time, and a peak with the same mass as oxidized cellulonodin-2 also emerged in the 22 h sample. We suspected that the minor products with the same mass as cellulonodin-2 and oxidized cellulonodin-2 were unthreaded variants of the peptides.^46^ To distinguish threaded and unthreaded variants, we treated the 22 h sample with a mixture of carboxypeptidases B and Y.^47^ Carboxypeptidases completely abolished peaks labelled with asterisks, suggesting that these two peaks corresponded to unthreaded species (Figure 4a). In contrast, lihuanodin remained stable and threaded when incubated at room temperature (Figure 4b).

**Figure 4:**
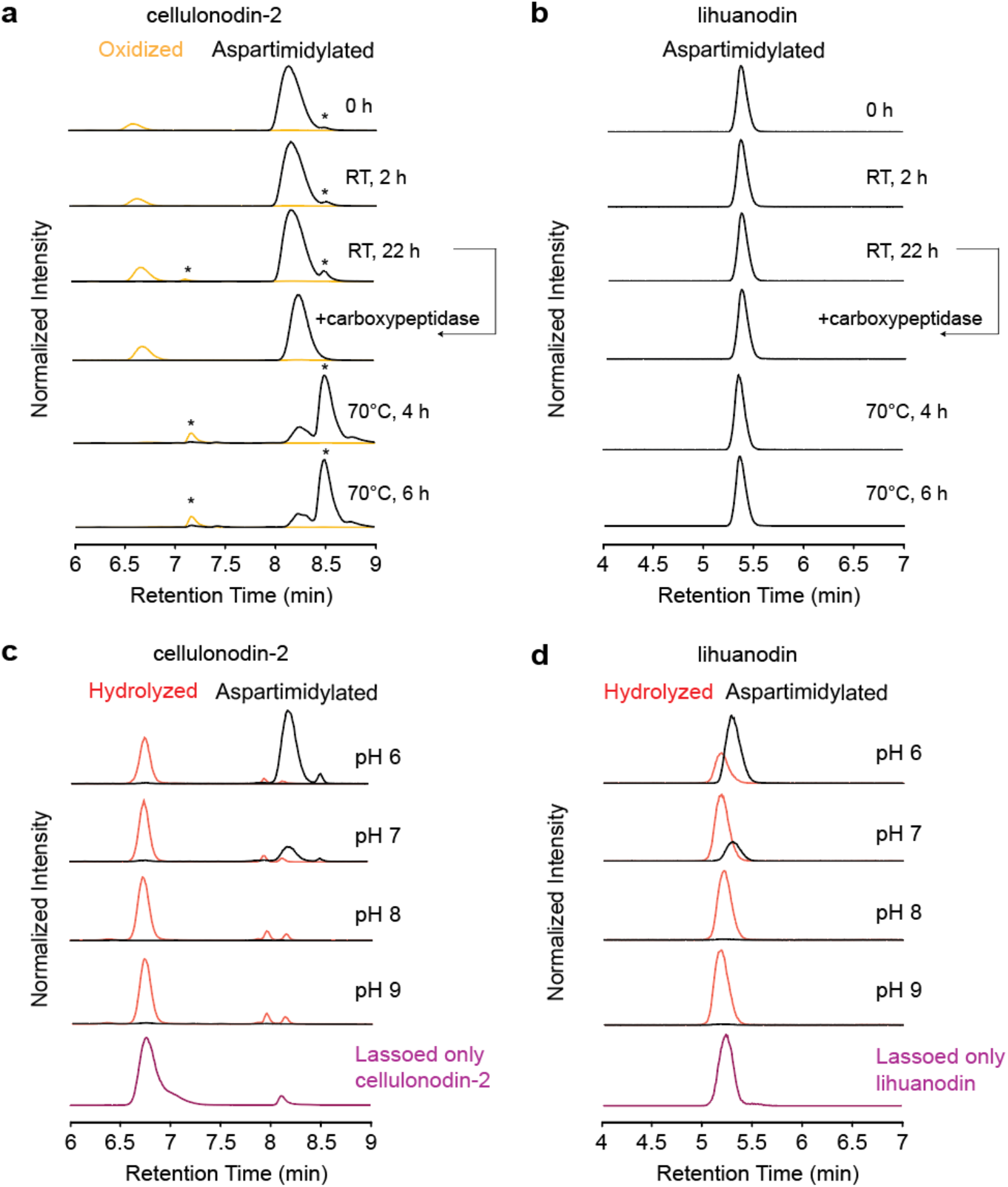
Stability Analysis of Cellulonodin-2 and Lihuanodin. Purified peptides were analyzed at 0 h, 2 h, and 22 h after incubation in ultrapure water at room temperature. The 22 h sample was further treated with carboxypeptidases. Putative unthreaded species are labelled with asterisks. For the heating assay at 70 °C, samples were withdrawn at 4 h and 6 h. Extracted ion current (EIC) traces for dehydrated, oxidized, and hydrolyzed species are labelled black, yellow, and red, respectively. **a)** Stability analysis of cellulonodin-2 in water at room temperature and 70 °C. The aspartimide moiety persists under both conditions, but a fraction of the peptide unthreads, especially at 70 °C. **b)** Stability analysis of lihuanodin in water at room temperature and 70 °C. The peptide remains threaded and the aspartimide remains intact under all conditions tested. **c)** Hydrolysis of cellulonodin-2 over 21 hours in 50 mM Tris-HCl buffer at pH = 6, 7, 8 and 9. Lassoed only cellulonodin-2 is shown for comparison (purple) and has the same retention time as the hydrolysis product. **d)** Hydrolysis of lihuanodin over 21 hours in 50 mM Tris-HCl buffer at pH = 6, 7, 8 and 9. Lassoed only lihuanodin (purple) has the same retention time as the hydrolysis product.

We also heated cellulonodin-2 and lihuanodin to 70 °C in ultrapure water, and analyzed the samples at 4 h and 6 h by LC-MS (Figure 4a, b). Most of cellulonodin-2 became the unthreaded variant in both 4 h and 6 h samples, indicating that higher temperature could speed up the unthreading process (Figure 4a). On the other hand, lihuanodin remained stable upon heating (Figure 4b). Remarkably, the aspartimide moieties in cellulonodin-2 and lihuanodin remained stable, even at 70 °C. Since the optimal growth temperature for *T. cellulosilytica* and *L. thermophila* is 45 °C, this result suggests that the aspartimide moiety remains stable under physiologically-relevant temperatures.

Succinimides are known to be susceptible to hydrolysis under alkaline conditions.^43^ Therefore, we incubated both cellulonodin-2 and lihuanodin in 50 mM Tris-HCl buffer at pH 6, 7, 8, and 9 over 21 h at 20 °C. In the presence of this higher ionic strength buffer, the hydrolysis of aspartimide was observed under all of these conditions, and the rate of hydrolysis was faster at higher pH for both peptides (Figure 4c, d). Surprisingly, cellulonodin-2 and lihuanodin were converted into only one hydrolysis product each. These products have the same retention times as the lassoed only peptides. This suggests that the succinimide in these peptides is opened regioselectively to give Asp and essentially no isoAsp. This is in stark contrast to the succinimides formed by canonical PIMTs, which open to give a mixture of products, typically 3:1 isoAsp to Asp.^48^

### NMR Structure of Lihuanodin

To further confirm that both cellulonodin-2 and lihuanodin contain an aspartimide moiety, we sought to solve NMR structures of these two peptides. We acquired TOCSY spectra of cellulonodin-2 in various conditions, including in 95:5 H_2_O:D_2_O at 4 °C and 20 °C, and in CD_3_OH at −20 °C, as well as TOCSY and NOESY spectra of lassoed only cellulonodin-2 in 95:5 H_2_O:D_2_O at 4 °C (Figure S10, S11). However, due to the unthreading behavior of cellulonodin-2, peaks on the spectra are either broad or indicate multiple conformers, making it infeasible to solve its structure. Encouraged by the stability of lihuanodin, we acquired TOCSY (60 ms mixing time) and NOESY (300 ms mixing time) spectra of lihuanodin in 95:5 H_2_O:D_2_O at 20 °C (Figure S12). The amide proton of T7 was missing on both TOCSY and NOESY spectra, suggesting the aspartimide moiety was indeed between the side chain of D6 and the backbone amide of T7 (Table S2). We calculated distance constraints based on the NOESY spectrum. Using CYANA 2.1, structural calculations revealed that all of the 20 top structures of lihuanodin contained a right-handed lasso, with a 9 aa isopeptide-bonded ring, a 4 aa loop, and a short 2 aa tail (Figure 5a, PDB: 7LCW). Extensive NOE cross-peaks between Y13/R14 and the ring residues were observed (Table S3), suggesting Y13 and R14 are the steric locks of the peptide. The side chain of W15 points away from the side chain of R14, indicating R14 and W15 can potentially serve as a branched dual lower lock (Figure 5b). Notably, the NOE cross-peaks strongly suggest that the thread is closer to the aspartimide side of the ring. This is reflected in the models built by CYANA in which the thread has an average distance of 4.2 ± 0.4 Å (Figure 5c).

**Figure 5:**
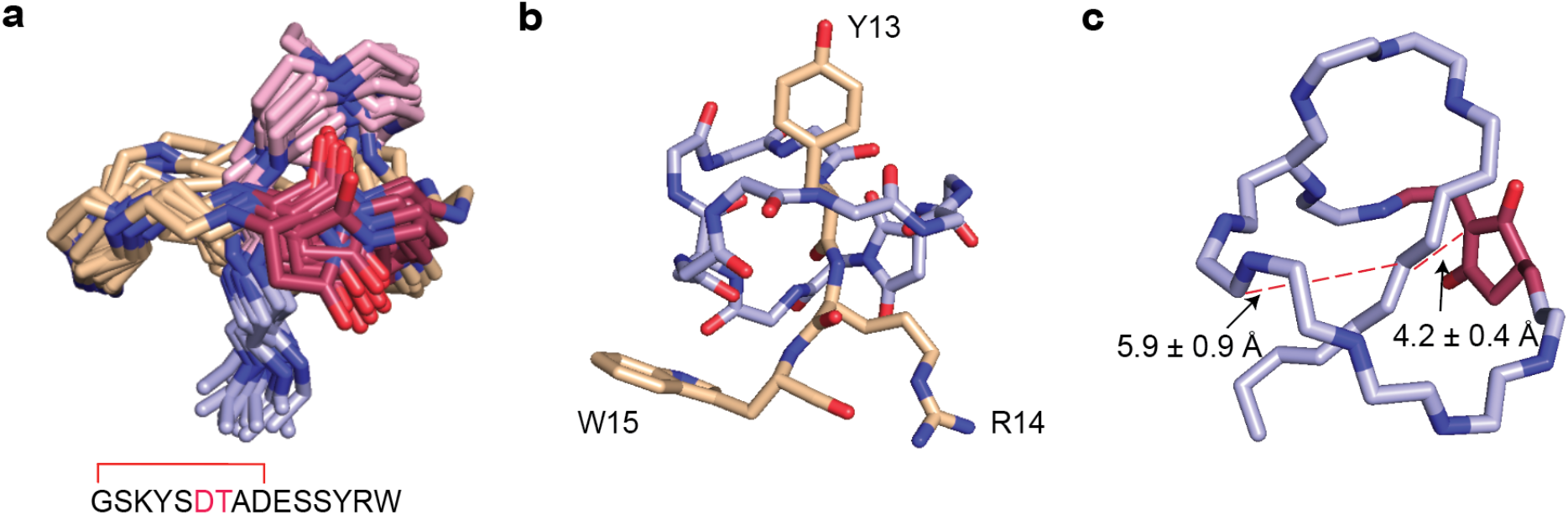
NMR Structure of Lihuanodin. **a)** Overlay of the top 20 structures of lihuanodin. Backbone carbon atoms of the ring is shown in wheat color, the loop is in pink, and the tail is in light blue. Backbone nitrogen atoms are colored royal blue throughout, and the aspartimide is shown in red. The sequence of lihuanodin is given below the structure showing the location of the isopeptide bond and the aspartimide (red). **b)** Top structure of lihuanodin, with the side chains of steric lock residues Y13, R14, and W15 shown. **c)** The C-terminal thread of lihuanodin is biased toward the aspartimide residue in the ring. The average distance measurement of the thread to the two sides of the ring is shown. The errors represent the standard deviation of the distance in all 20 structures.

### Mutagenesis of Cellulonodin-2 and Lihuanodin Core Peptides

A hallmark of RiPPs biosynthetic machinery is its tolerance to different substrate sequences. Therefore, we performed extensive mutagenesis on cellulonodin-2 and lihuanodin to illustrate the plasticity of the biosynthetic enzymes as well as the promiscuity of the associated methyltransferases, with a focus on the conserved DTAD motif in the ring. Our NMR data on lihuanodin showed that the aspartimide forms between the side chain of D6 and the backbone of T7. In the case of cellulonodin-2, amino acid substitutions to D6 and T7 led to an accumulation of lassoed only and methylated species except in the case of T7S, suggesting that the hydroxyl group (-OH) of the Thr or Ser side chain can aid in the aspartimide formation (Figure 6a). To further validate our experimental results, calculations at the B3LYP-D3(BJ)/def2-TZVP^49,50^ level of density functional theory were performed for a series of model compounds with the structure AcAsp(OCH_3_)XxxNHCH_3_ (Xxx = Thr, Val, Ser, and Ala). Further details about these DFT experiments can be found in the Supporting Information. Initial calculations showed lower gas-phase proton affinities of the model dipeptides’ amide groups for Xxx = Thr and Ser compared to corresponding non-polar residues with a similar steric demand, Val and Ala (Figure S13). Furthermore, we calculated p*K*_a_ value differences (Δp*K*_a_) using the conductor-like polarizable continuum model and a proton exchange thermodynamic cycle. Again, the more polar Thr and Ser give lower p*K*_a_ values than their non-polar counterparts indicating that Thr and Ser can assist in aspartimide formation (Figure S13). This calculation helped us validate why T7 was highly conserved in precursor sequences. However, it is worth noting that the *n*+1 amide’s p*K*_a_ would also be affected by the conformational freedom of a peptide sequence, which is certainly restricted in a lasso peptide.^51^

**Figure 6:**
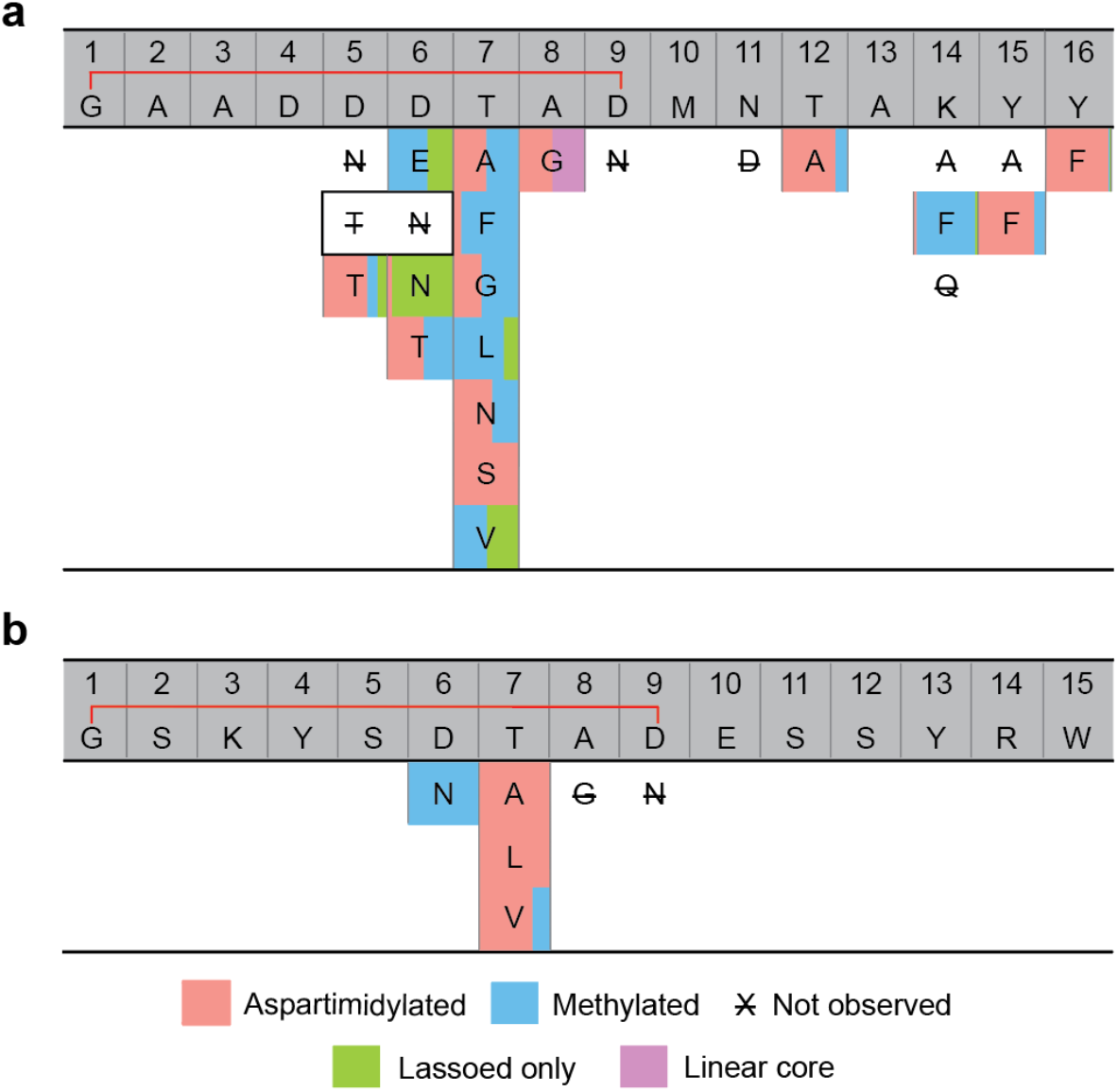
Effects of Single Amino Acid Substitutions of Cellulonodin-2 and Lihuanodin on Lasso Formation and Aspartimide Formation. Each of the indicated variants were heterologously expressed under the same conditions as the wild-type peptides. The bracket indicates the isopeptide bond between G1 and D9 for both peptides. The blocks for each variant are colored according to the proportion of each species observed in LC-MS. The residues enclosed in the same border indicate a D5T D6N double substitution. The percentage of each species is reported in **Figure S14. a)** Cellulonodin-2 variants. **b)** Lihuanodin variants. For both peptides, the D6N substitution reduces aspartimide formation. Substitutions to T7, the residue contributing its amide to aspartimide, have varying effects, but all variants are at least still methylated.

When T7 in cellulonodin-2 was changed to residues with non-polar side chains (V, L, F) by mutagenesis, we observed a lower percentage of the dehydrated product compared to amino acids with small or polar side chains (A, N), indicating a bulky side chain on the *n*+1 residue also negatively affects aspartimide formation. In addition, in the case of D6T and D6E, we observed a methylated lassoed species to be the major product, indicating that TceM has promiscuous activity against another D residue in the ring when D6 is substituted. We further constructed the A8G variant, and the major products are linear core and the dehydrated lasso, suggesting A8 may be crucial for TceC recognition and isopeptide bond formation (Figure 6a, Figure S13). As expected, D9N resulted in no lasso production, as the ability for isopeptide bond formation was abolished. We also mutated the DTAD motif in lihuanodin to test whether both peptides behave similarly (Figure 6b). The D6N variant completely abolished the additional dehydration, suggesting D6 was indeed the site of the modification. In contrast to what we have seen in the case of cellulonodin-2, lihuanodin T7A still allows lihuanodin to be fully dehydrated. As a result, we constructed lihuanodin T7V and T7L variants. Both of these variants were still effectively dehydrated, suggesting that the lihuanodin ring is more readily methylated or dehydrated than the cellulonodin-2 ring. The lihuanodin A8G variant was not produced, suggesting that it is not a substrate for the lasso peptide biosynthetic machinery, in agreement with the results for cellulonodin-2. As expected, the D9N variant abolished lasso formation, as it did for cellulonodin-2.

Since we could not directly observe the aspartimide moiety in cellulonodin-2 by NMR, we sought to reaffirm that the nucleophilic side chains were not involved in bond formation via mutagenesis. We mutated all residues with nucleophilic side chains in the loop and tail regions. T12A, Y15F, and Y16F resulted in predominantly the dehydrated lassoed peptides, indicating that they were not involved in the bond formation (Figure 6a). In contrast, the major product of the K14F variant was the methylated product, likely due to the fact that F is bulkier than K and could potentially obstruct the aspartimide formation in the ring. Unfortunately, we were unable to verify this hypothesis, as the more conservative variant K14Q did not express (Figure 6a, Figure S14).

### Role of the C-Terminal Domain of TceM and LihM

Since both TceM and LihM are predicted to be PIMT homologs, we performed an amino acid sequence alignment of TceM, LihM, and canonical PIMTs from several bacteria and eukaryotes. TceM and LihM consist of an N-terminal region that is homologous to PIMTs (∼220 aa), but also contain a unique C-terminal domain of ∼110–130 aa (Figure S15).^52^ We first constructed a truncated TceM by removing the entire C-terminal domain, and coexpressed it with the lassoed only cellulonodin-2 plasmid. The C-terminal truncation of TceM led to no methylation or aspartimidylation on the lassoed only cellulonodin-2 (Figure S16), suggesting the C-terminal domain was required for its activity.

Subjecting a selection of sequences of PIMT homologs associated with lasso peptide clusters to Multiple EM for Motif Elicitation (MEME) revealed a conserved WXXXGXP motif in the C-terminal domain (Figure 7a).^53^ To evaluate the importance of these residues, we constructed TceM W323A and P329A as well as LihM W306A and P312A. The W323A substitution on TceM nearly abolished the aspartimide formation for cellulonodin-2, with 96.5% (by LC-MS peak area) of the variants remaining lassoed only (Figure 7b). TceM P329A also led to an increase in the lassoed only species compared to the wild-type TceM (18% vs 0.5% by LC-MS peak area, Figure 7b). On the other hand, both LihM W306A and P312A only slightly increased the percentage of the lassoed only and methylated species compared to the wild-type (Figure 7c). Instead, for the dehydrated species, two peaks in the case of LihM W306A, or a broad peak in the case of P312A were observed, indicating these residues were important for the proper maturation of lihuanodin.

**Figure 7:**
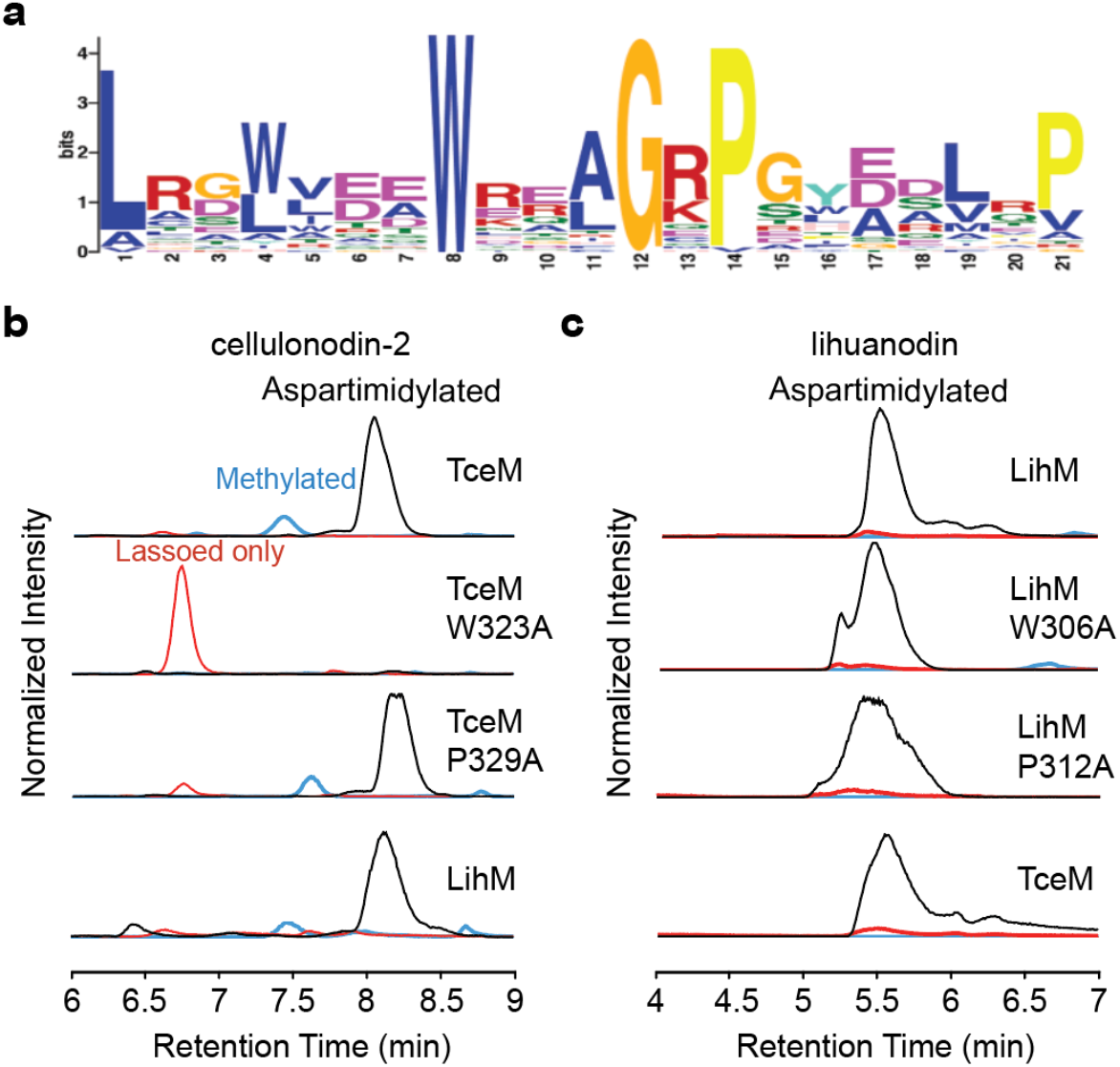
Role of a Conserved Motif in the C-terminal Domain of TceM and LihM. **a)** Sequence logo of a conserved C-terminal motif of methyltransferases associated with lasso peptide BGCs. **b)** From top to bottom, lassoed only cellulonodin-2 coexpressed with WT TceM and TceM W323A, P329A and LihM. Black traces are lassoed and aspartimidylated, blue traces are lassoed and methylated, and red traces are lassoed only. The W323A substitution disrupts methylation severely. **c)** From top to bottom, lassoed only lihuanodin coexpressed with WT LihM and LihM W306A, P312A and TceM. Color code is the same as part b. Both variants of LihM are competent at methylation, in contrast to the cellulonodin-2 case. However, both the W306A and P312A LihM lead to broader product peaks, suggesting that multiple products may be formed.

Considering the possibility that the C-terminal segment in TceM and LihM could be responsible for recognizing the DTAD motif in the lasso peptide ring, we sought to investigate whether TceM and LihM had promiscuous activity on their non-cognate substrates. Under our optimized heterologous expression conditions for both peptides, TceM and LihM can successfully form the aspartimide in lassoed only lihuanodin and cellulonodin-2, respectively, to an extent similar to its cognate methyltransferase (Figure 7b, c). Since the amino acid sequences of cellulonodin-2 and lihuanodin and the two methyltransferases are divergent other than the DTAD motif in the lasso and the WXXXGXP motif in the enzyme, the result strongly suggests that the conserved WXXXGXP motif in methyltransferases plays a role in substrate recognition.

### Reconstitution of TceM and LihM Activities *in vitro*

Previous examples of enzymes that carry out PTMs on lasso peptides have been shown to modify the loop and tail regions of the peptide, and recognized only the linear precursor as the substrate.^28,31,34^ Since cellulonodin-2 and lihuanodin are the first examples of lasso peptides in which ring residues are post-translationally modified, we sought to identify the minimal substrates for TceM and LihM by reconstituting their activities *in vitro*. Both TceM-His_6_ and LihM-His_6_ were expressed and purified using nickel affinity chromatography to homogeneity at good yields (2.5 and 4.8 mg/L culture respectively, Figure S17). We first tested lassoed only cellulonodin-2 and lihuanodin as substrates for TceM and LihM respectively, and used 10 equiv. of SAM as the methyl donor and analyzed the reaction products using LC-MS. In 30 min, for cellulonodin-2, all starting material was reacted, with a product distribution of 24% methylated, and 76% dehydrated (Figure 8a). For lihuanodin, the starting material was nearly exclusively converted to the dehydrated product (Figure 8d). We repeated the experiment in the absence of SAM, and observed no modifications in both cases (Figure S18), indicating SAM was the co-substrate for both methyltransferases as expected. In order to monitor the progression of the reaction, we took time points of both *in vitro* reactions (Figure 8a, d). In the case of cellulonodin-2, starting material was all methylated in 5 min, and the methylated species was gradually converted to the dehydrated species over time (Figure 8a). A similar pattern was also observed in the case of lihuanodin, with all starting material getting methylated in 5 min. However, the aspartimide formation step proceeded faster in the case of lihuanodin compared to cellulonodin-2. The *in vitro* results also agreed with the *in vivo* results that a higher percentage of dehydrated products could be made in the case of lihuanodin compared to cellulonodin-2. Increasing reaction time from 30 min to 2 h did not change the product distributions (Figure S19).

**Figure 8:**
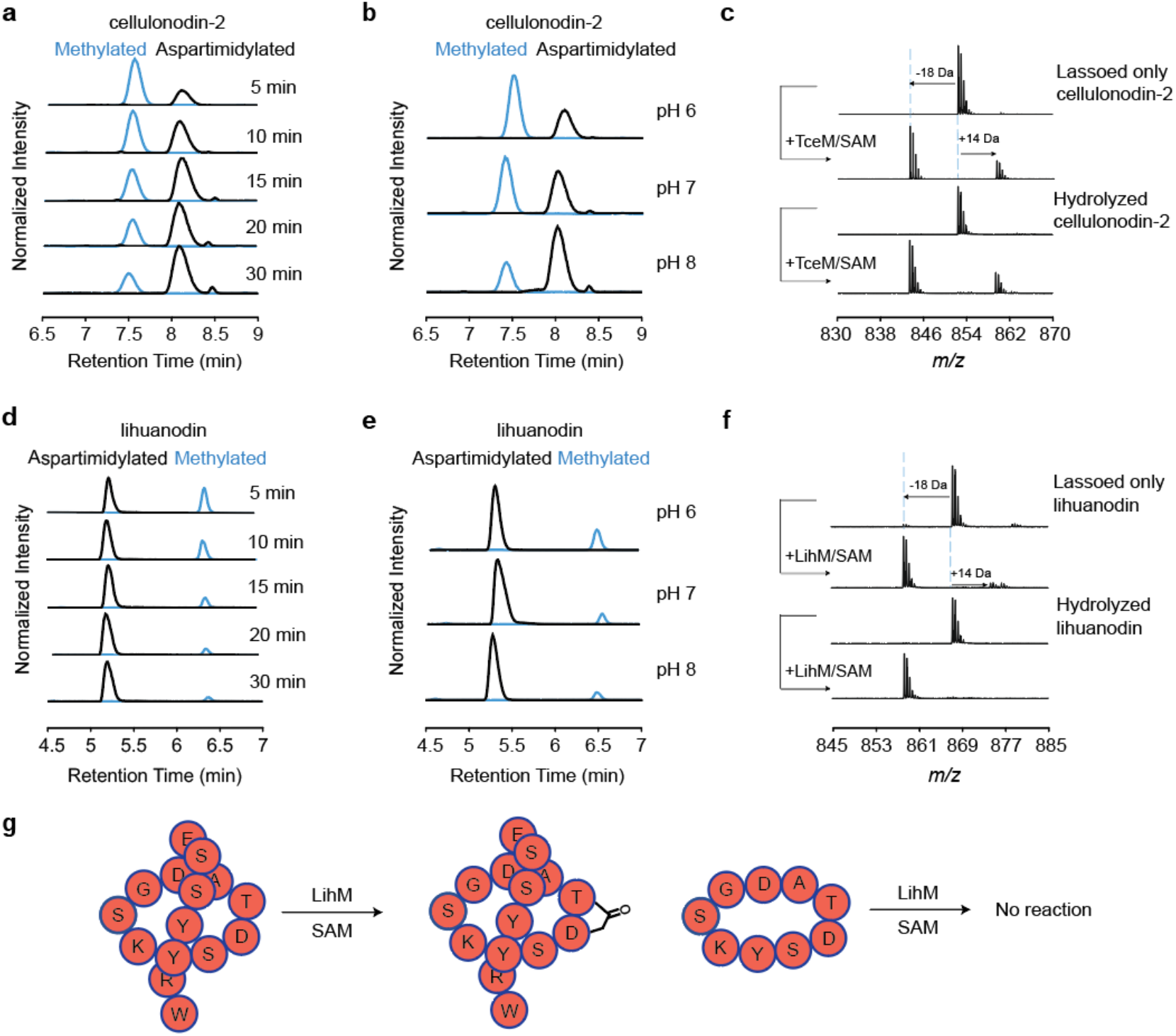
*In vitro* Maturation of Cellulonodin-2 and Lihuanodin. **a)** Time course of lassoed only cellulonodin-2 reacted with TceM. Reactions contained 100 μM lasso peptide substrate, 1μM enzyme, 1 mM SAM in 50 mM Tris-HCl, pH 7.4 and were incubated at 37 °C. Under these conditions, all starting material is converted to the methylated form within 5 min. Aspartimide formation lags behind methylation. **b)** Lassoed only cellulonodin-2 reacted with TceM at varying pH as indicated. Reaction conditions were otherwise as in part a); reactions were carried out for 30 min. As expected, aspartimide formation is more rapid at higher pH. **c)** Mass spectra of lassoed only and hydrolyzed cellulonodin-2 reacted with TceM for 30 min at pH 7.4; concentrations as in part a. The hydrolyzed cellulonodin-2 sample was generated by incubating cellulonodin-2 (with the aspartimide moiety) in Tris-HCl, pH 8 overnight. In both cases, the starting material is completely consumed and a mixture of methylated and dehydrated products are observed. This suggests that the hydrolyzed material is identical to lassoed only cellulonodin-2. **d)** Time course of lassoed only lihuanodin reacted with LihM. Conditions as described in part a. Like the cellulonodin-2 case, all starting material is consumed within 5 min, but the aspartimide formation is more rapid, being nearly complete after 30 min. **e)** Lassoed only lihuanodin reacted with LihM at varying pH; conditions as in part b. Again, higher pH accelerates aspartimide formation. **f)** Mass spectra of lassoed only and hydrolyzed lihuanodin reacted with LihM. Like the cellulonodin-2 case in part c, hydrolyzed lihuanodin acts similarly to lassoed only lihuanodin, suggesting that these species are identical. **g)** LihM recognizes lassoed only lihuanodin as its substrate, but not its isopeptide-bonded ring alone (Figure S19).

We then sought to determine whether the lasso peptide structure of cellulonodin-2 and lihuanodin was required for modification by TceM and LihM, respectively. We first tested the linear precursor and linear core of cellulonodin-2 as substrates for TceM, and saw no modifications in both cases (Figure S20). Moreover, we constructed the cellulonodin-2 ring and lihuanodin ring using solid phase peptide synthesis (SPPS),^54^ and also observed no modifications using them as substrates (Figure S20). These results suggested that the entire lasso structure was required for enzyme recognition (Figure 8g). Since LihM and TceM were able to modify lassoed only cellulonodin-2 and lihuanodin *in vivo* respectively, we further carried out these two reactions *in vitro* and observed the two methyltransferases could indeed be used interchangeably to mature both cellulonodin-2 and lihuanodin (Figure S21).

We further carried out the *in vitro* reactions for both cellulonodin-2 and lihuanodin in 50 mM Tris-HCl buffers at pH 6, 7, and 8, and observed a higher percentage of the dehydrated species at higher pHs, agreeing with the fact that the backbone amide of *n*+1 residue would be more nucleophilic under basic conditions (Figure 8b, e). Since both host organisms of cellulonodin-2 and lihuanodin are thermophiles with optimal growth at 45 °C, we also performed *in vitro* reactions at 50 °C and observed a similar extent of reaction at the elevated temperature (Figure S22).

Additionally, we performed *in vitro* reactions using hydrolyzed cellulonodin-2 and lihuanodin (Figure 4) as substrates. The cognate methyltransferases could re-dehydrate the hydrolyzed products in 30 min (Figure 8c, f), suggesting the aspartimide-containing cellulonodin-2 and lihuanodin are the intended natural products in the native hosts. As we noted above, it appears that the aspartimides in cellulonodin-2 and lihuanodin are both regioselectively hydrolyzed to Asp. These *in vitro* results affirm that assertion; the hydrolysis products can be fully converted back to the aspartimidylated lasso peptides.

## Discussion

In this work we have described a novel post-translational modification for RiPPs, aspartimide, in the ring region of two lasso peptides, cellulonodin-2 and lihuanodin, both from thermophilic Gram-positive bacteria. The BGCs for these lasso peptides include an *O*-methyltransferase that installs the aspartimide in a manner reminiscent of the activity of protein L-isoaspartyl methyltransferases (PIMTs). Whereas canonical PIMTs form aspartimide from isoaspartate as an intermediate in protein backbone repair, the PIMTs associated with these lasso peptide BGCs instead function on aspartate. The modified aspartate is the first amino acid in a highly conserved tetrapeptide (DTAD). Neither the linear lasso peptide core peptide nor a macrocyclic peptide corresponding to the lasso peptide ring are substrates for this enzyme; only the lasso peptide functions as a substrate (Figure S20). The aspartimide moiety in these peptides is noteworthy because of its stability; in low ionic strength environments, the aspartimide resists hydrolysis. *In vitro* experiments show that, when the aspartimide in these lasso peptides hydrolyzes, it does so in a regioselective fashion, opening up only to aspartate (Figure 4). This substrate is identical to the starting material, allowing for further rounds of conversion to the aspartimide by the PIMT. This line of evidence points to the conclusion that aspartimide-modified lasso peptides are the intended final product of this pathway. While aspartimides have notoriety in the solid-phase peptide synthesis field and in pharmaceutical processing as undesired side products,^44,55^ the PIMT homologs funnel substrate toward aspartimide products in this enzymatic transformation. Another PIMT homolog colocalized with a lanthipeptide BGC was reported by van der Donk and colleagues.^52^ In contrast to our results here, the final lanthipeptide product was hydrolyzed, and consisted of a mixture of Asp and isoAsp residues in the peptide backbone.

Besides its altered substrate specificity (i.e. modifying Asp instead of isoAsp), the PIMT homologs associated with lasso peptide BGCs have several unique features. Both of the enzymes studied here, TceM and LihM, are exceptionally fast methyl transfer catalysts, at least with respect to other published reports of PIMTs. In our *in vitro* time course studies of the conversion of lasso peptides to their methylated forms (Figure 8), all of the substrate was methylated within 5 minutes. This means that the minimum specific activity for these enzymes is 516 nmol•min^−1^•mg protein^−1^ for TceM and 521 nmol•min^−1^•mg protein^−1^ for LihM. These activities compare favorably with the specific activity measurements reported for the *P. furiosus* PIMT (164 nmol•min^−1^•mg protein^−1^ at pH 4 and 68 °C) and human PIMT (11.4 nmol•min^−1^•mg protein^−1^ at pH 7 and 37 °C).^56^ Another feature of TceM and LihM that differentiates them from canonical PIMTs is the presence of an “extra” domain of ∼110–130 aa at the C-terminus of these enzymes. Our work here shows that this C-terminal domain is required for TceM activity (Figure S16). Furthermore, substitution of a conserved tryptophan residue within this C-terminal domain is deleterious to the catalytic activity of both TceM and LihM (Figure 7). Homology modeling using iTasser^57^ and structure prediction using HHPred^58^ of this C-terminal domain did not lead to any clues about its structure or function (Figure S3). We hypothesize that the C-terminal domain is involved in substrate recognition of the relatively bulky lasso peptide substrate, perhaps positioning it for efficient catalysis at the methyl transfer site of the enzyme. Further structural studies on TceM and LihM are expected to shed light on the remaining puzzles of this enzyme’s function: how do TceM and LihM recognize a specific Asp only within the lasso structure and methylate it so quickly?

The aspartimide modification, which rigidifies the peptide backbone, is reminiscent of other backbone rigidification chemistries, such as oxazol(in)es and thiazol(in)es, which appear both in RiPPs and in nonribosomal peptides.^59,60^ Because of its electrophilicity, the aspartimide is expected to be much less stable than these moieties. The aspartimides in cellulonodin-2 and lihuanodin exhibit some stability, even at elevated temperatures (Figure 4), in low ionic strength environments. The reasons for this relative stability are likely related to the lasso structure in which the aspartimide is embedded. The aspartimide is part of 9 aa macrocycle, which limits torsion angles available to the peptide backbone. The fact that when these peptides hydrolyze they do so in a regiospecific manner suggests that perhaps the lasso peptide structure hinders the attack of water on one side of the succinimide.

While we have not directly studied the physiological role of the aspartimide here, our data suggests some potential scenarios. While our data clearly show that the final product of the PIMT homologs is the aspartimide, the persistence of this functional group will be dependent on the final disposition of the lasso peptide. For example, if these peptides remain intracellular, we expect them to remain in the aspartimide form as long as SAM is available and the PIMT homolog is expressed. However, if these peptides are secreted from their producers into a basic environment, such as the warm manure pile from which *T. cellulosilytica* was isolated,^61^ the lasso peptide will revert to its aspartate form. However, if the peptide were to be secreted into a more neutral, dilute environment, such as pond water, we expect the aspartimide to persist for some time. Of note is that the lihuanodin producer, *L. thermophila*, grows best on dilute media with no added NaCl.^62^ We noticed that, in *E. coli*, production of lassoed only cellulonodin-2 (i.e. without the aspartimide) led to decreased cell growth relative to the production of cellulonodin-2 (Figure S23). This result opens up the possibility that lassoed only cellulonodin-2 functions as an antibiotic in the *E. coli* cytoplasm, and the aspartimide moiety could be functioning as a resistance mechanism in the native producer. Another scenario, especially if the aspartimide persists for a long period of time in its environment, is that cellulonodin-2 and/or lihuanodin could be functioning as suicide inhibitors to a receptor or enzyme. For example, a key lysine residue on such proteins could attack the aspartimide, leading to irreversible formation of an amide bond. In summary, we have presented a new PTM for RiPPs which is installed by remarkably fast methyltransferase enzymes with unusual substrate specificity. Future studies will be directed both at characterization of these enzymes as well as understanding the role of the aspartimide in the bioactivity of these lasso peptides.

## Acknowledgements

We thank I. Pelczer (Princeton University NMR Facility) for help with acquiring NMR spectra. This work was supported by National Institutes of Health Grant GM107036 and a grant from Princeton University School of Engineering and Applied Sciences (Focused Research Team on Precision Antibiotics). L.C. was supported by an NSF Graduate Research Fellowship Program under Grant DGE-1656466. J.D.K. was supported in part by training grant T32 GM7388. H.V.S. was supported by the Deutsche Forschungsgemeinschaft (DFG Research Fellowship, GZ: SCHR 1659/1-1).

