## Supplementary Information for "Cellulonodin-2 and Lihuanodin: Lasso Peptides with an Aspartimide Post-translational Modification"

#### Supplementary Materials and Methods

##### Materials and Equipment

*Thermobifida cellulosilytica* (DSM 44535) was obtained from the German Collection of Microorganisms and Cell Cultures (DSMZ), and genomic DNA was isolated using the Qiagen DNeasy Blood and Tissue Kit standard protocol. Genomic DNA of *Lihuaxuella thermophila* (DSM 46701) was ordered directly from DSMZ. *Escherichia coli* XL-1 Blue were used for all recombinant DNA procedures. All primers were ordered from Integrated DNA Technologies (IDT). Restriction enzymes, Q5 DNA polymerase, and T4 DNA ligase were purchased from New England Biolabs (NEB). The sequences of all primers used are provided in Table S1. All sequencing results were confirmed by Sanger sequencing (Genewiz). Acetonitrile was purchased from Sigma-Aldrich. Semi-preparative reverse-phase HPLC was performed using a Zorbax 300SB-C18 (9.4 mm x 250 mm, 5 μm) column with an Agilent 1200 series instrument. LC-MS and LC-MS/MS experiments were done using a Zorbax 300SB-C18 (2.1 mm x 50 mm, 3.5 μm) column with a 1260 Infinity II system coupled to an Agilent 6530 qTOF instrument. NMR experiments were done using a Bruker Avance III 800 MHz instrument.

#### Cloning

##### Cellulonodin-2 gene cluster

The gene cluster containing *tceACB1B2M* was amplified from the genomic DNA and was digested using *Xba*I and *Hind*III restriction sites and ligated into pASK-75 cloning vector. The GTG start codon of B2 was changed to ATG using overlap PCR. The intergenic non-coding region between *tceA* and *tceC* was removed by digesting the vector with *Xba*I and *Blp*I, as the region encodes has a predicted hairpin on the RNA level that may inhibit translation (Figure S23). The resulting plasmid is pMB4. The gene cluster was also refactored into a pQE-80 vector, with the *tceA* gene placed under an inducible-T5 promoter and the *tceCB1B2M* genes placed under a constitutive *mcjBCD* promoter from the microcin J25 gene cluster. Briefly, *tceA* was first cloned into pQE-80 using *Eco*RI and *Hind*III restriction sites, creating plasmid pMB2. The *tceCB1B2M* genes were then PCR-amplified from pMB4. The constitutive promoter of *mcjBCD* from the microcin J25 gene cluster was appended upstream of *tceCB1B2M* in an overlap PCR. This PCR product was then cloned in the forward direction into pMB2 using *Nhe*I and *Nco*I restriction sites, forming pMB3.

Cellulonodin-2 variants were cloned using pMB3 as a template, and *tce*A was swapped with mutated precursor using *Xho*I and *Hin*dIII/*Nhe*I restriction sites. Plasmids either eliminating the *tceM* (pLC13) or with a C-terminally truncated *tceM* gene in pQE80 were cloned using pMB4 as a template, and *tceCB1B2* was placed between *Nhe*I and *Nco*I restriction sites instead.

The *tceM* gene was amplified from the genomic DNA and appended with a C-terminal 6x histidine tag. The PCR product was digested using *Nco*I and *Hind*III restriction sites and ligated into the first multiple cloning site of pRSF duet cloning vector, forming pJDK127. Genes encoding TceM variants were cloned using pJDK127 as a template using overlap PCR, and ligated to the pRSF Duet vector between the *Nco*I and *Hind*III restriction sites.

##### Lihuanodin gene cluster

The gene cluster containing *lihMACB1B2* was amplified from the genomic DNA and was digested using *Xba*I and *Sal*I restriction sites and ligated into the pASK-75 vector. The resulting plasmid is called pMO7. The gene cluster was also refactored into a coexpression system. In the pQE-80 vector, the *lihA* gene was placed under an inducible T5 promoter and the *tceCB1B2* gene placed under a constitutively-expressed *mcjBCD* promoter from the microcin J25 gene cluster. Briefly, *lihA* was first cloned into pQE-80 using *Eco*RI and *Hin*dIII restriction sites, creating plasmid pMO8. The *lihCB1B2* operon was then PCR-amplified from pMO7. The start codon of *lihC* was changed from TTG to ATG. The constitutive promoter of *mcjBCD* from the microcin J25 gene

cluster was appended upstream of *lihCB1B2* in an overlap PCR. This PCR product was then cloned in the forward direction into pMO8 using *Nhe*I and *Nco*I restriction sites, forming pMO9. In the pRSF-Duet cloning vector, the *lihM* gene was placed under an inducible T7 promoter on its own, creating plasmid pLC30. Lihuanodin variants were cloned using pMO9 as a template, and *lihA* was swapped with mutated precursor using *Xho*I and *NheI* restriction sites.
*LihM* was amplified from the genomic DNA and appended with a C-terminal 6x Histidine tag. The PCR product was digested using *Eco*RI and *Sal*I restriction sites and ligated into the multiple cloning site of pQE80, creating pLC29. LihM truncation and mutants in pRSF-Duet construct was cloned using pLC30 as a template, and ligated between *Nco*I and *Hind*III in pRSF Duet vector.

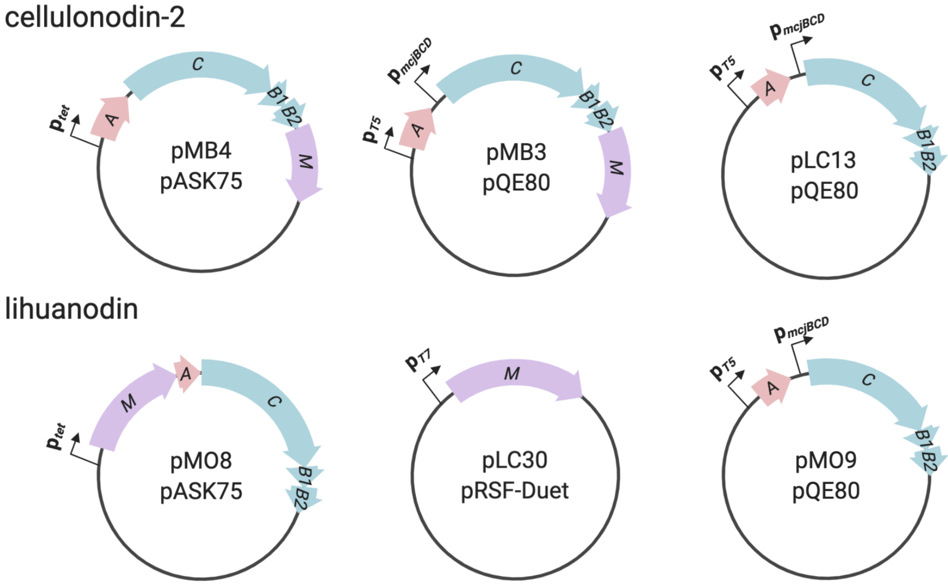

#### Expression and Purification of Cellulonodin-2, Lihuanodin, and their variants

Expression of cellulonodin-2, lihuanodin, as well as their variants (point mutants and lassoed only cellulonodin-2 and lihuanodin) were carried out using *E. coli* BL21 in M9 media supplemented with 0.00005 wt% thiamine, 1 mM of $MgSO_4$, 20 amino acids (0.05 g/L of each amino acid), and 100 mg/L ampicillin (for lihuanodin, also with 50 mg/L kanamycin). The typical expression scale performed was a 500 mL culture in a 2L flask. For expression, a 5 mL overnight *E. coli* BL21 culture was grown in LB with 100 mg/L ampicillin (for lihuanodin, also with 50 mg/L kanamycin) and then subcultured into the supplemented M9 media to a starting $OD_{600}$ of 0.02. The M9 cultures were then grown at 37 °C, 250 rpm until an $OD_{600}$ of 0.2. The cultures were then induced with 200 µg/L anhydrotetracycline (aTc) if they contained a pASK75-based plasmid, or 1 mM of isopropyl-ß-D-thiogalactopyranoside (IPTG) if they contained a pQE-80-based or a pRSF-duet based plasmid. For cellulonodin-2 and its variants, expression was carried out at 37 °C, 250 rpm for 4 hours. For lihuanodin and its variants, expression was carried out at 37 °C, 250 rpm for 2 days. Cells were harvested by pelleting the cells at 4000 x *g*, 4 °C for 20 minutes. The cell pellet was washed with 1X PBS. After removing the PBS, 10 mL/L of culture of 100% methanol was added along with glass beads. The pellet was then vortexed until homogeneous. The mixture was spun down at 8000 x g and the supernatant was collected and dried using a rotovap. The dried samples were resuspended in 100% water (5000x compared to expression volume), spun down, and used for HPLC and LC-qTOF analysis.

Typically for RP-HPLC purification, 60-80 µL of sample was injected onto the semi-preparative column. The mobile phase A consisted of water with 0.1% trifluoroacetic acid and phase B consisted of acetonitrile with 0.1% trifluoroacetic acid. The gradient program used was as follows: 10% B from 0-1 minute, linear gradient from 10-45% B from 1-20 minutes, linear gradient from 45-90% B from 20-25 minutes, and isocratic elution at 90% B from 25-27 minutes.

For LC-MS and LC-MS/MS analysis, 3-10 µL of purified cellulonodin-2, lihuanodin, and their variants or 15 uL of TceM and LihM were injected onto the Zorbax 300SB-C18 (2.1 mm x 50 mm, 3.5 µm, for peptides) column or the XBridge Protein BEH C4 column (2.1mm x 50 mm, 300 Å, for proteins) with an Agilent 1260 Infinity II system for separation. The mobile phase A consisted of water with 0.1% formic acid, and phase B consisted of acetonitrile with 0.1% formic acid. The gradient used was: 5% B from 0-1 minute, linear gradient from 5-45% B from 1-20 minutes, linear gradient from 45-90% B from 20-25 minutes, and isocratic elution at 90% B from 25-30 minutes. The separated species were directly sprayed to an Agilent 6530 q-TOF instrument for mass detection and fragmentation.

#### Expression and Purification of TceM-His$_6$ and LihM-His$_6$

The plasmid containing the C-terminal 6xHis tagged TceM was transformed into *E. coli* BL21 DE3 Δ*slyD* cells and grown up overnight in LB with 50 mg/L kanamycin. The cells were subcultured at an $OD_{600}$ = 0.02 into 500 mL of LB with 50 mg/L kanamycin and allowed to grow at 37 °C. Similarly, the plasmid containing the C-terminal 6xHis tagged LihM was transformed into *E. coli* BL21 Δ*slyD* cells and grown up overnight in LB with 100 mg/L ampicillin. The cells were subcultured at an $OD_{600}$ = 0.02 into 500 mL of LB with 100 mg/L ampicillin and allowed to grow at 37 °C. Once the $OD_{600}$ = 0.5, expression was induced using 1 mM IPTG, also at 37 °C. The cells were grown for 4 hours before being spun down at 4000 x *g*. The pellet from a 500 mL culture was resuspended in 10 mL of 50 mM $NaH_2PO_4$, 300 mM NaCl, 10 mM imidazole, pH 8.0 before freezing at -80 °C. The resuspended cell pellet was thawed in ice water and 10 mg of lysozyme was added, followed by incubation at 4 °C for 20 minutes. The sample was then sonicated on ice to lyse the cells and spun down at 4000 x *g* for 10 minutes at 4 °C. The supernatant was removed and spun down an additional time at 8000 x *g* for 10 minutes. The clarified lysate was then incubated with 1 mL of Ni-NTA resin (Qiagen) while rotating for 1 hour. The mixture was added to an empty gravity column, and the flowthrough was passed over the resin an additional time. The resin was then washed with 10 mL of 50 mM $NaH_2PO_4$, 300 mM NaCl, 20 mM imidazole pH 8.0 and twice with 10 mL of 50 mM $NaH_2PO_4$, 300 mM NaCl, 50 mM imidazole pH 8.0. The protein was then eluted with 50 mM $NaH_2PO_4$, 300 mM NaCl, 250 mM imidazole pH 8.0. The samples were run on SDS-PAGE and the most concentrated samples were pooled and buffer exchanged using a PD-10 desalting column into 50 mM Tris-HCl, pH 7.4. Purified and buffer exchanged TceM and LihM were frozen in 10% glycerol and stored in aliquots to conserve activity.

#### Carboxypeptidase Assay

Lasso peptides (5 µL) were digested with 1 U carboxypeptidase B (Sigma-Aldrich) and 1 U carboxypeptidase Y (Affymetrix) in 50 mM sodium acetate, pH = 6.0, with a total volume of 50 µL for 16 hours at 20 °C. In a typical experiment, 5-10 µL of the digested mixture was injected onto LC-MS for analysis.

#### Aspartimide Hydrolysis Assay

Solutions of cellulonodin-2 and lihuanodin (100 µM) were incubated in 50 mM of Tris-HCl buffer at pH 6, 7, 8, and 9 for a total volume of 50 µL for 21 hours at 20 °C. The incubated peptides (10 µL) were injected onto LC-MS for analysis.

#### Hydrazine Reaction

In the reaction mixture, cellulonodin-2 and lihuanodin were added to organic solvents (acetonitrile for cellulonodin-2 and DMSO for lihuanodin since lihuanodin is not soluble in acetonitrile) to a final concentration of 0.1 mM and hydrazine (from a solution of 35% hydrazine in water, Sigma-Aldrich) was added to a final concentration of 2 M, with a final volume of 50 µL. The reaction mixture was incubated at room temperature for 30 mins before getting analyzed by LC-MS.

#### Structural Elucidation of Lihuanodin

TOCSY and NOESY NMR spectra (Figure S12) were acquired at the Princeton University Department of Chemistry NMR Facilities on an A8 Avance III HD 800-MHz NMR spectrometer (Bruker) with a triple-resonance cryoprobe. The NMR sample of lihuanodin was prepared in 95/5 $H_2O/D_2O$ at a concentration of 1.6 mM. A list of proton shifts and long-range NOE correlations are listed in Supplemental Tables 2-3.

#### NMR Structure Calculations of Lihuanodin

TOCSY (60 ms mixing time) and NOESY (300 ms mixing time) spectra of lihuanodin were acquired in 95/5 $H_2O/D_2O$ at 293 K. Cross-peak positions and volumes in this spectrum were manually picked and assigned in MestReNova. These were given as input data for the calculations, which were performed in CYANA 2.1 on a Linux cluster. We incorporated explicit distance constraints of 1.4 Å for the isopeptide bond between the N of Gly1 and the $C_\gamma$ of Asp9, and the succinimide linkage between the $C_\gamma$ of Asp6 and the N of Thr7. These distances were based on the structure of capistruin (PDB 6N61)[1] and lysozyme (PDB 1AT5).[2] Seven cycles of combined NOESY assignment and structure calculation were performed, and the top 20 structures were generated. The generated structures were further relaxed using MMFF94 forcefield for energy minimization.[3] The calculated conformers were visualized in PyMOL.

#### Reconstitution of TceM and LihM Activities *in vitro*

The *in vitro* assays were comprised of 1 mM of *S*-adenosylmethionine (SAM), 100 µM of lassoed only cellulonodin-2 or lihuanodin, 1 µM of TceM or LihM in 50 mM Tris-HCl buffer at pH 7.4, with a final reaction volume of 50 µL. The reactions were set up either at 37 °C or 50 °C. The reaction mixtures were incubated at various time points, and subsequently sampled and quenched using 5 µL 10% formic acid. The samples were then analyzed using LC-MS.

#### *E. coli* BL21 *Δpcm* Construction

*E. coli* BL21 *Δpcm::kan* was generated by P1 transduction from the Keio collection, where the donor strain was E. coli K12 MG1655 *Δpcm::kan. kan* signifies kanamycin-resistance marker. The recipient strain *E. coli* BL21 *Δpcm::kan* was confirmed by colony PCR.

#### Solid Phase Peptide Synthesis

Peptides were synthesized manually on a 0.25 mM scale using Rink amide resins (Chem-Impex International Inc).The Fmoc group was deprotected using 20% piperidine (Alfa Aesar)/*N,N*-Dimethylformamide (DMF, Alfa Aesar), for 20 min to obtain a deprotected peptide-resin. Asp(*O*-allyl)OH amino acid (1.25 mM, Ambeed Inc.) was loaded on the Rink resin first. Fmoc-protected amino acids (0.75 mM, AnaSpec Inc) were sequentially coupled on the resin using *N,N,N',N'*-Tetramethyl-*O*-(1H-benzotriazol-1-yl)uranium hexafluorophosphate (HBTU, 0.75 mM, Chem-Impex International Inc.) and *N,N*-Diisopropylethylamine (DIEA, 0.75 mM, Tokyo Chemical Industrial Co. Ltd.) for 2 h at room temperature. To facilitate the isopeptide bond formation between Gly1 and Asp9 side chain, a solution of Tetrakis(triphenylphosphine)palladium(0), $Pd(PPh_3)_4$ (20 mg, Sigma-Aldrich), phenylsilane (72 µL, Sigma-Aldrich) in dichloromethane (DCM, 3 mL, Honeywell), and the peptide was left on a shaker for 40 minutes. The resin was washed with DCM (3 X 2 min). The above reaction was repeated, and then the resin was washed with DCM (3 X 2 min), MeOH (3 X 2 min), and DMF (3 X 2 min). The palladium catalyst was removed from the resin by washing it with DIEA (2 X 2 min), followed by washing with DMF (2 X 2 min).

Next, the peptide was treated with 20% piperidine/DMF to remove the Fmoc protecting group. Macrocyclization was achieved by treating the resin with *N,N′*-diisopropylcarbodiimide (DIC, Alfa Aesar), 1-Hydroxy-7-azabenzotriazole (HOAt, Chemscene) , and a catalytic amount of 4-Dimethylaminopyridine (DMAP, Alfa Aesar) in DMF on a shaker overnight. The solution was drained, and the resin was washed with DMF. To further confirm formation of cyclic peptides, the resin was cleaved using a cocktail of 95:2.5:2.5 trifluoroacetic acid (TFA, Millipore):triisopropyl silane (TIPS, Tokyo Chemical Industrial Co. Ltd.):water (v/v/v) for 2 h. The resin was then removed by filtration, and the resulting solution was evaporated. Cyclic peptides were purified by semi-preparative chromatography and analyzed by LC-MS.

#### Supplementary Information for Refactoring Cellulonodin-2 and Lihuanodin Biosynthetic Gene Clusters (BGCs)

In order to check for production of a lasso product from the lasso peptide BGC from *T. cellulosilytica,* the cluster was cloned and inserted into a pASK75 vector under an anhydrotetracycline (aTc) inducible promoter. The native ORF that encoded *tceB2* had a GTG start codon and was changed to ATG in order to improve expression for *E. coli* BL21. In addition, the native BGC contained a 126 base pair intergenic region between the stop codon of *tceA* and the start codon of *tceC.* The putative RNA transcript of this intergenic region was input into *RNAStructure*,[4] a server that predicted secondary structure of nucleic acids. The predicted secondary structure of transcribed RNA in fact showed multiple possible hairpin formations, which could make it difficult for ribosome to traverse the transcription (Figure S24). As a result, the hairpin-forming intergenic region was replaced with a 25 base-pair sequence that would not give to any secondary structure. This cluster was further refactored into a pQE80 vector, where the precursor gene *tceA* was placed under an isopropyl-β-D-thiogalactopyranoside (IPTG) inducible promoter, and the biosynthetic enzymes and the methyltransferase were placed under the constitutive promoter native to the biosynthetic enzymes needed for microcin J25 synthesis (Figure 2a). Both constructs could lead to cellulonodin-2 production, while the pQE80 version resulted in a better yield by ~2-fold. The yield of cellulonodin-2 is still quite low at 30 μg/L.

Having successfully expressed cellulonodin-2 heterologously in *E. coli* BL21, we applied a similar refactoring strategy for the lasso peptide BGC from *L. thermophila*. The cluster was also cloned and inserted into a pASK75 vector under an anhydrotetracycline (aTc) inducible promoter. In addition, we further refactored the cluster into a co-expression system. In the pQE80 vector, the precursor gene *lihA* was placed an IPTG inducible protomer, and the biosynthetic enzymes were also placed under the constitutive promoter native to the biosynthetic enzymes needed for microcin J25 synthesis (Figure 2a). The start codon of *lihC* was changed from TTG to ATG. The associated methyltransferase was placed under a T7 promoter in the pRSF-Duet vector (Figure 2a). While we observed no product formation from the pASK75 construct, the coexpression system allowed the production of lihuanodin at 35 μg/L.

#### Supplementary Information on DFT Calculations of Dipeptide $AcAsp(OCH_3)XxxNHCH_3$

To find suitable starting structures, the conformational space of model dipeptide $AcAspThrNHCH_3$ was scanned by a molecular dynamics simulation in the CHARMM36[5] force field using an explicit water box under periodic boundary conditions. Several conformations were selected and energy-minimized in the MMFF94[3] force field. Afterwards, three conformations were further optimized using Grimme's HF-3c method[6] and the continuum solvation model CPCM[7] as implemented in ORCA[8] 4.2.1. For each state, three minimized conformations have been constructed, which show single point energies with small energy differences. Frequency calculations of the optimized conformations do not show any imaginary frequencies. The anionic forms were generated by removing the amide bond's proton and structures were re-optimized in gas phase and solution. For the anionic succinimide state, structure optimization led to a deprotonation of the adjacent amide bond by the formed methoxide molecule. The lowest-energy conformer of each state was picked and single point energy calculations were performed at the B3LYP-D3(BJ)/def2-TZVP[9,10] level of density functional theory using the RIJK approximation. The SCF Convergence tolerance was set to "TightSCF" and the CPCM module was used. For all calculations, the DFT-D3 dispersion correction with Becke-Johnson damping (D3BJ) has been applied.[11,12]

For transition state optimization, the nudged elastic band[13] (NEB) method as implemented in ORCA was used to find a minimum energy path between reactant and product state and the corresponding saddle point on the potential energy surface. The NEB-TS command was applied utilizing 10 images and three different convergence tolerance values for regular images, climbing images and saddle-point optimization (Max(|Fp|): $2\cdot10^{-2}$ Eh/Bohr and RMS(Fp): $1\cdot10^{-2}$ Eh/Bohr; Max(|Fp|): $2\cdot10^{-3}$ and RMS(F): $1\cdot10^{-3}$ Eh/Bohr; and Max(|Fp|): $3\cdot10^{-4}$ Eh/Bohr and RMS(F): $1\cdot10^{-4}$ Eh/Bohr; respectively). All derived transition states showed an imaginary frequency mode.

p$K_a$ values were calculate by using the proton exchange thermodynamic cycle shown in Figure S25. This cycle includes an explicit water molecule, which is considered to be less prone to errors compared to the direct cycle of deprotonation.[14] The p$K_a$ is calculated by equations 1–4.[15,16] The dissociation of an acid $AH_{aq}$ in aqueous solution is given by the equilibrium in eq. 1. The p$K_a$ can be directly calculated from the free energy change $\Delta G_{aq}$ in solution (eq. 2), which consist of the free gas phase reaction energy $\Delta G_g$ and the differences between solvation free energies $\Delta\Delta G_s$ (eq. 3 and 4). Solvation free energies have been calculated by using the CPCM module.

$$\mathrm{AH_{aq}} + \mathrm{H_2O_{aq}} \rightleftharpoons \mathrm{A^-_{aq}} + \mathrm{H_3O^+_{aq}} \qquad \text{(eq. 1)}$$

$$\mathrm{p}K_\mathrm{a} = \frac{\Delta G_\mathrm{aq}}{\ln(10)RT} - 1.74 \qquad \text{(eq. 2)}$$

$$\Delta G_\mathrm{aq} = \Delta G_\mathrm{g} + \Delta G_\mathrm{s}(\mathrm{A^-}) - \Delta G_\mathrm{s}(\mathrm{AH}) + \Delta G_\mathrm{s}(\mathrm{H_3O^+}) - \Delta G_\mathrm{s}(\mathrm{H_2O}) = \Delta G_\mathrm{g} + \Delta\Delta G_\mathrm{s} \qquad \text{(eq. 3)}$$

$$\Delta G_\mathrm{g} = G_\mathrm{g}(\mathrm{A^-}) - G_\mathrm{g}(\mathrm{AH}) + G_\mathrm{g}(\mathrm{H_3O^+}) - G_\mathrm{g}(\mathrm{H_2O}) \qquad \text{(eq. 4)}$$

#### Supplementary Figures

**a**

```
                         10        20        30        40
fuscanodin      ME-----KKKYTAPQLAKVGEFKEATGWYTAEWGLELIFVFPRFI
cellulonodin-1  MQEKKTQRKKYTRPQIAKAGGFKESTGWIQGKWGLEIYLIFPRYL
cellulonodin-2  MD-----EEIYQPPELVELGDAREVTLGAADDDTADMNTAKYY
```

**b**

| Protein | C | B1 | B2 |
|---|---|---|---|
| fuscanodin vs. cellulonodin-1 | 77% | 72% | 88% |
| fuscanodin vs. cellulonodin-2 | 26% | 37% | 33% |
| cellulonodin-1 vs. cellulonodin-2 | 26% | 39% | 34% |

**Figure S1: Comparison of Lasso Peptide Gene Cluster in *T. fusca* and the Lasso Peptide Gene Clusters in *T. cellulosilytica*. a)** Sequence alignment of the precursors of these three lasso clusters. Leader sequence is shown in black, isopeptide bonded ring in blue, and the thread in grey. Glu is at the point of cyclization and is colored red. The putative steric lock residues are labeled orange. Note the similarity between fuscanodin and cellulonodin-1, whereas the cellulonodin-2 sequence differs significantly. **b)** Sequence identity between maturation enzymes is high for fuscanodin and cellulonodin-1. The maturation enzymes for cellulonodin-2 have much lower sequence similarity to those of the other two peptides.

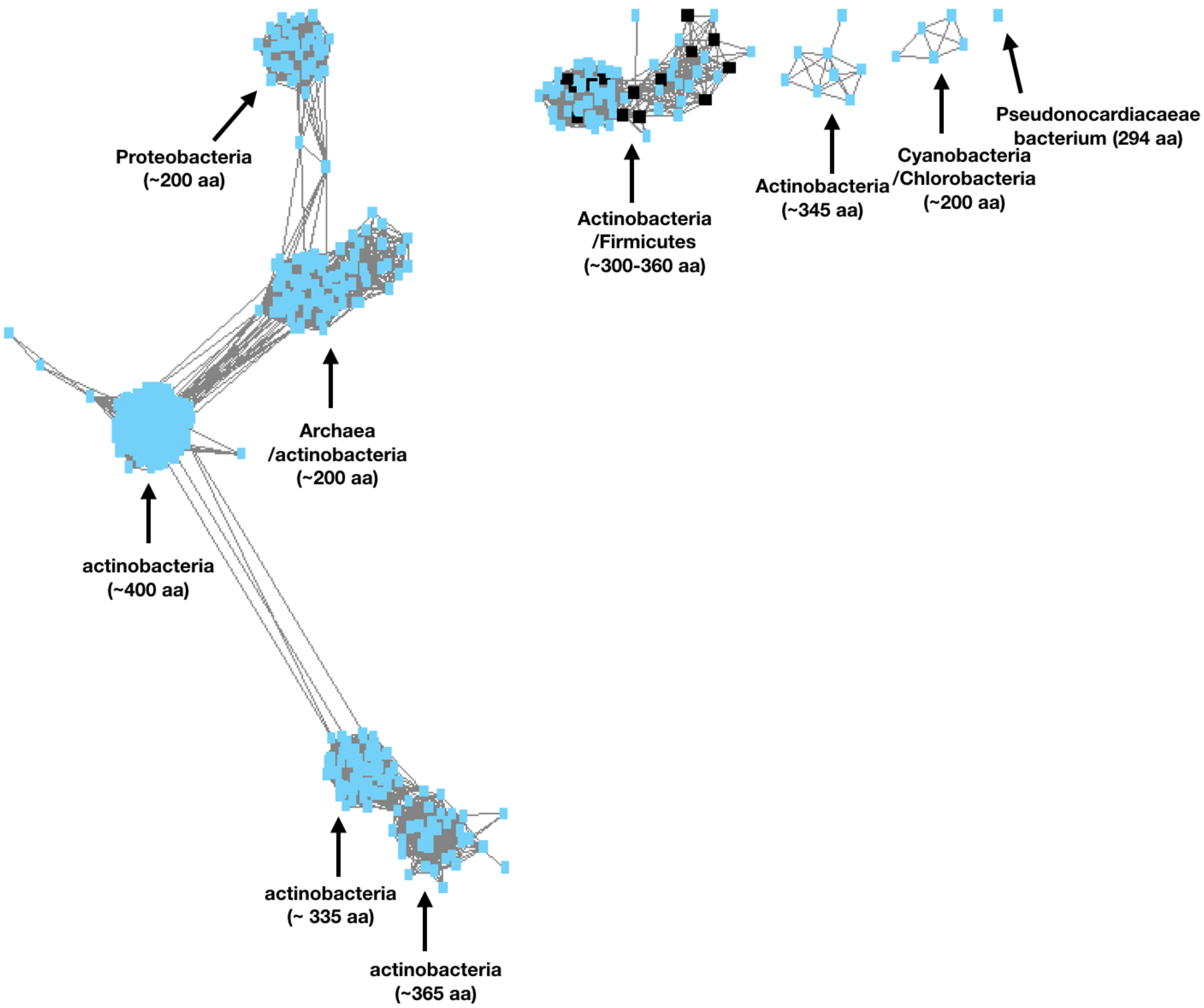

**Figure S2: Sequence Similarity Network (SSN) of TceM.** Using the cellulonodin-2 methyltransferase TceM as a query, 1000 methyltransferase sequences were surveyed and the SSN was generated using EFI-EST.[17] A threshold value of 40 was used to generate the SSN. Many proteins in this SSN were annotated as Protein-L-isoaspartyl *O*-methyltransferase (PIMT). All PIMT homologs in this SSN associated with lasso peptides clustered together (black squares). Of the 48 lasso-associated methyltransferases found by BLAST (Figure S4), 14 were found in this SSN cluster, including LihM.

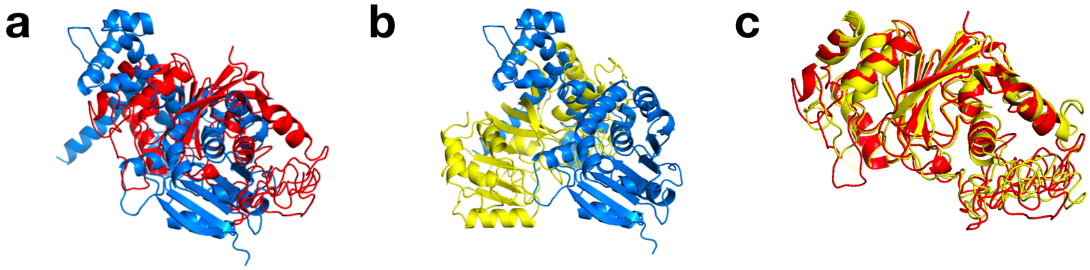

**Figure S3: Comparison of Homology Models of O-Methyltransferases in Lasso Peptide BGCs.** Using iTasser, homology models were generated for TceM, LihM, and StspM, the methyltransferase from lassomycin-like gene clusters. StspM methylates the C-terminus of its cognate lasso peptide. Alignment of StspM and TceM/LihM showed minimal homology. Predicted structure of StspM, TceM and LihM were colored in blue, red, and yellow, respectively. **a)** Structural alignment of StspM (blue) and TceM (red). **b)** Structural alignment of StspM (blue) and LihM (yellow). **c)** Structural alignment of TceM (red) and LihM (yellow). The disordered region at the bottom of the structure corresponds to the C-terminus of TceM and LihM.

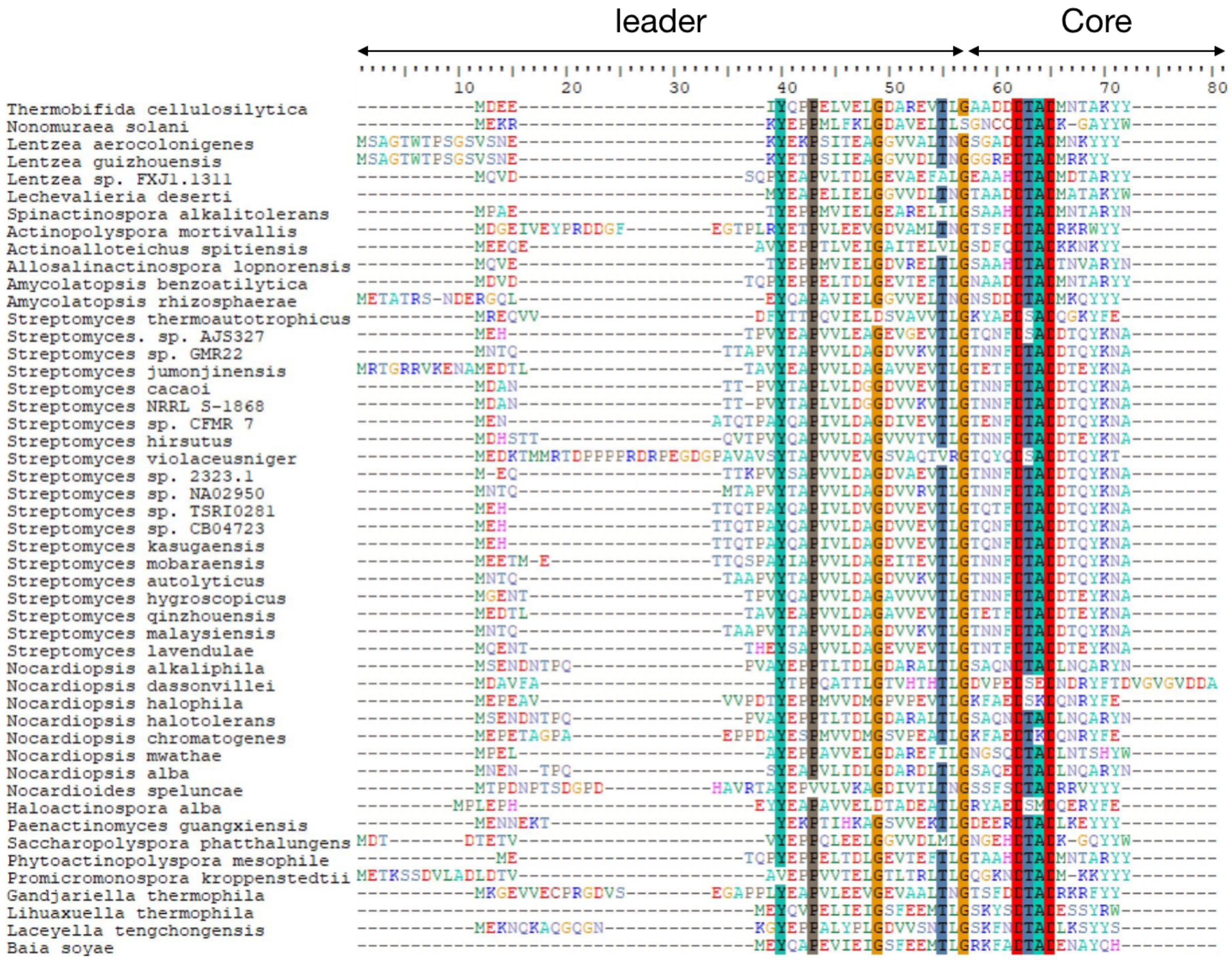

**Figure S4: Sequence Alignment of All Putative Lasso Peptide Precursors from BGCs that Contain a PIMT homolog.** The top 43 organisms are actinobacteria, while the bottom 5 organisms are firmicutes. The leader and core region of the precursor are indicated. In the core peptide region, there is a conserved G residue at position 1, and a conserved DTAD tetrapeptide at position 6-9.

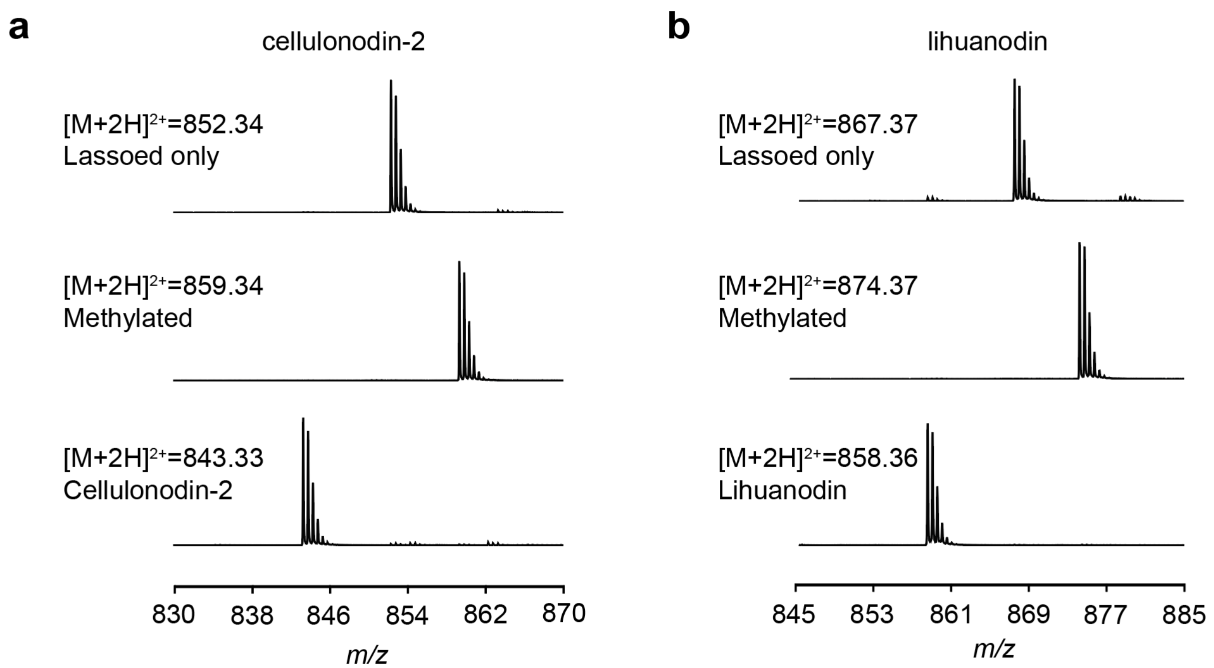

**Figure S5: Mass Spectra of Species Observed in Cellulonodin-2 and Lihuanodin BGC Expression. a)** Mass spectra of lassoed only cellulonodin-2 (1702.68 Da), methylated lassoed cellulonodin-2 (1716.69 Da) and cellulonodin-2 (1684.66 Da). **b)** Mass spectra of lassoed only lihuanodin (1732.74 Da), methylated lassoed lihuanodin (1746.75 Da) and lihuanodin (1714.72 Da).

a

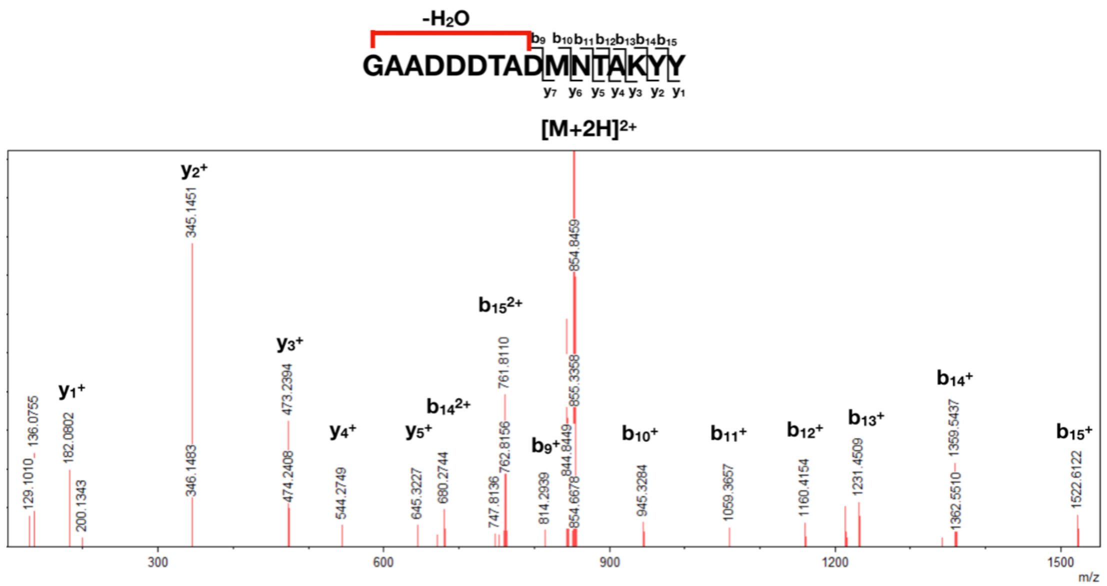

b

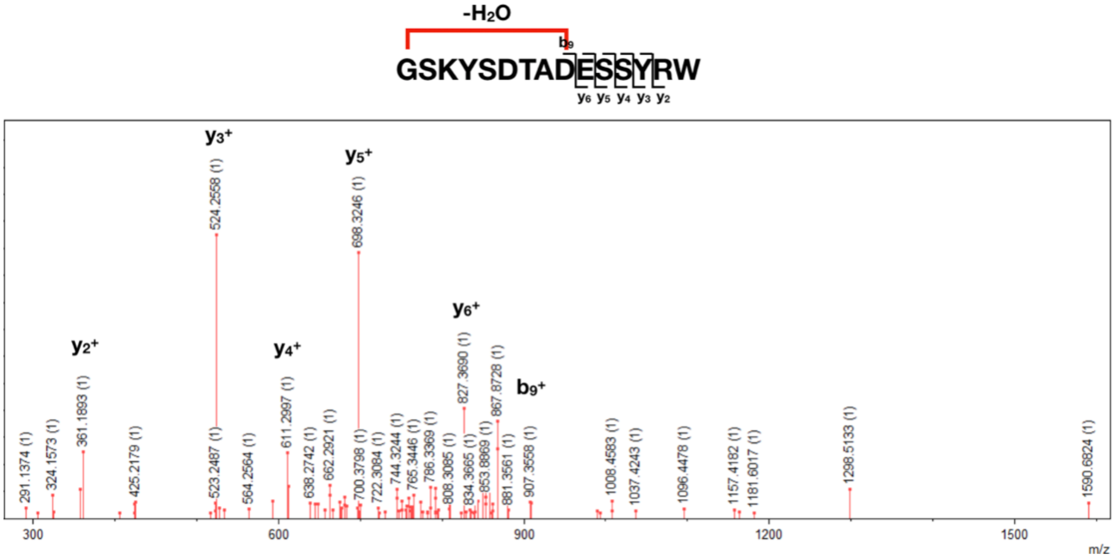

**Figure S6: MS/MS Fragmentation of Lassoed Only Cellulonodin-2 and Lihuanodin**. The ring fragment contained only one dehydration, indicating that TceM/LihM was responsible for the second dehydration on each of the peptide *in vivo*. **a)** MS/MS fragmentation of lassoed only cellulonodin-2. b) MS/MS fragmentation of lassoed only lihuanodin.

**a**

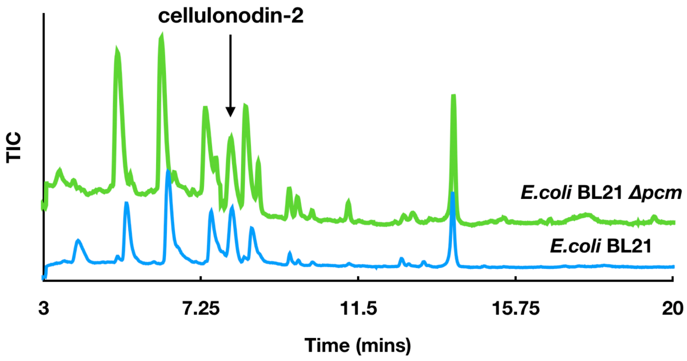

**b**

| | Aspartimidy -lated lasso | Methylated lasso | Lassoed only | Intensity of the major product |
|---|---|---|---|---|
| ***E. coli*** **BL21** | 84.8% | 14.2% | 1.0% | $3.1 \times 10^7$ |
| ***E. coli*** **BL21 *Δpcm*** | 83.6% | 14.1% | 2.3% | $2.4 \times 10^7$ |

**Figure S7: Comparison of Cellulonodin-2 expression using *E. coli* BL21 and *E. coli* BL21 *Δpcm*.** The *pcm* gene in *E. coli* encodes for a canonical PIMT. **a)** LC-MS traces. **b)** product distributions using two different strains. Deletion of *pcm* does not influence the final product distribution in these heterologous expression experiments.

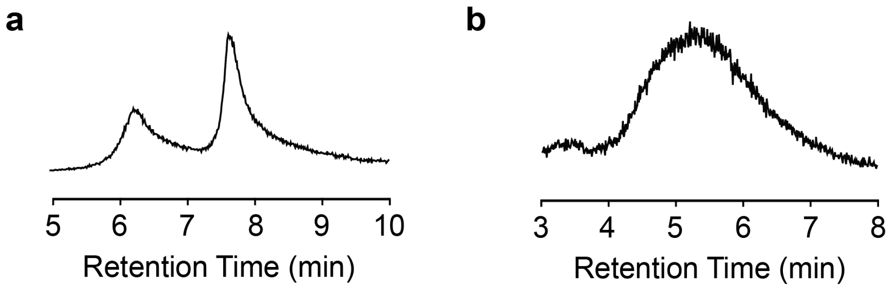

**Figure S8: Extracted Ion Current Chromatograms (EICs) of Cellulonodin-2 and Lihuanodin Reacted with Hydrazine. a)** EIC of cellulonodin-2 with hydrazide addition**. b)** EIC of lihuanodin with hydrazide addition**.** Both cellulonodin-2 and lihuanodin are fully reacted at room temperature for 30 mins in ACN (cellulonodin-2) and in DMSO (lihuanodin).

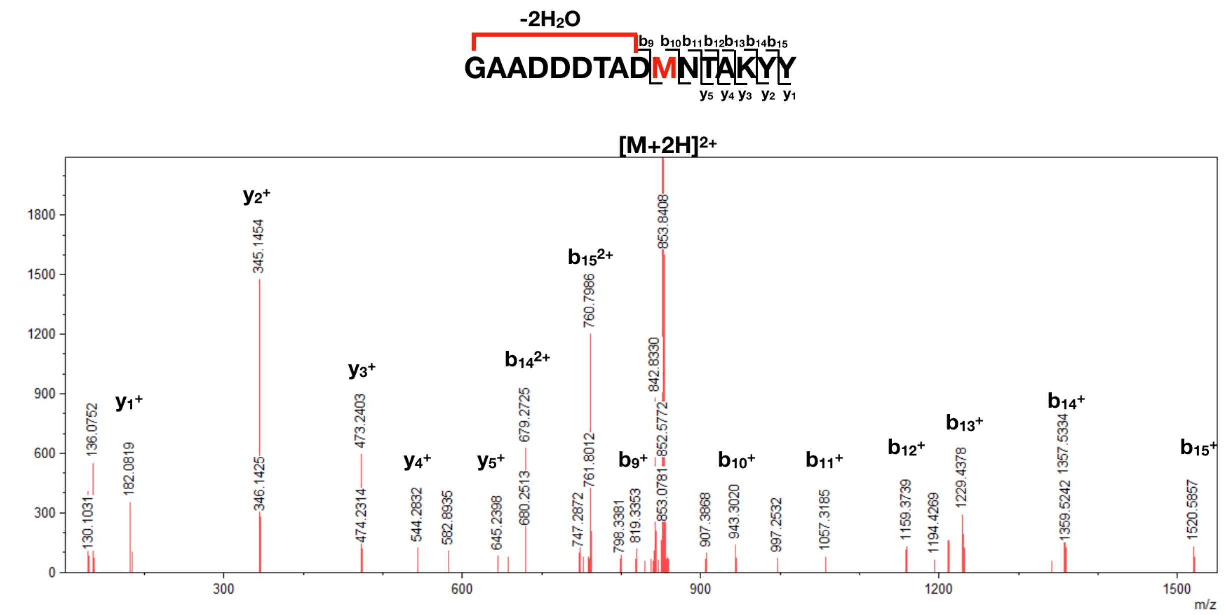

**Figure S9: MS/MS fragmentation of oxidized cellulonodin-2.** The Met that gets oxidized is labelled red. The fragmentation pattern supports that the oxidation occurs on M10.

**a**

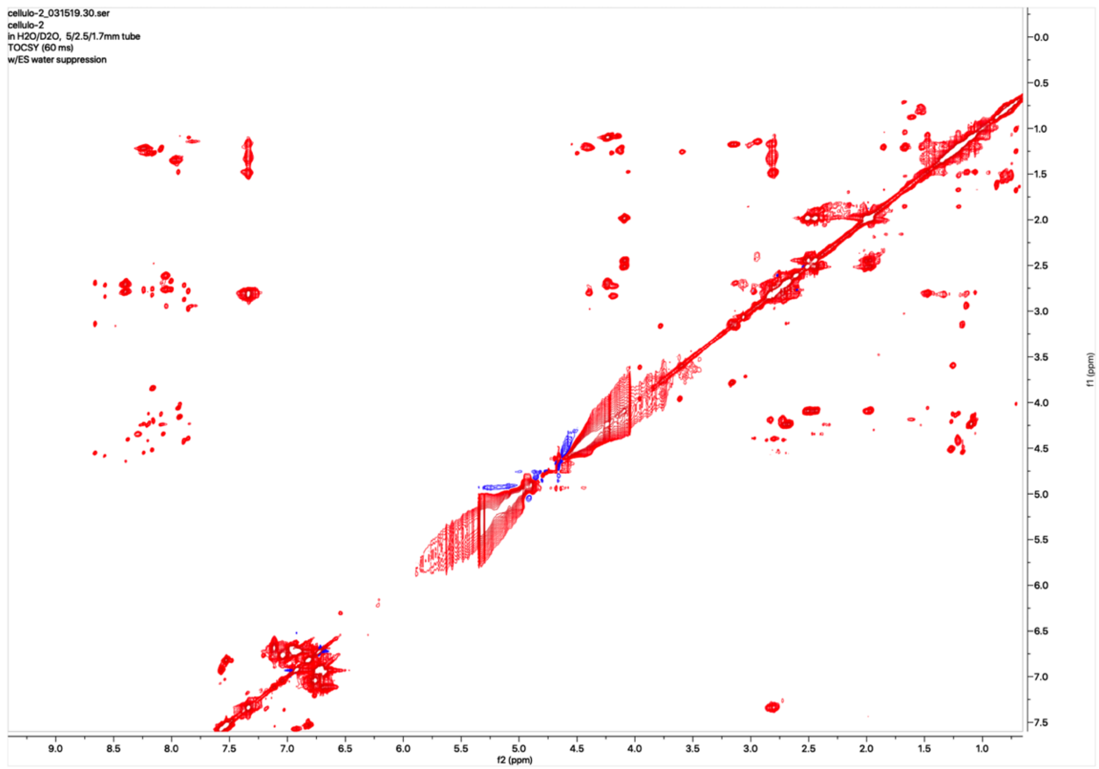
cellulo-2_031519.30.ser
cellulo-2
in H2O/D2O, 5/2.5/1.7mm tube
TOCSY (60 ms)
w/ES water suppression
f2 (ppm)
f1 (ppm)

**b**

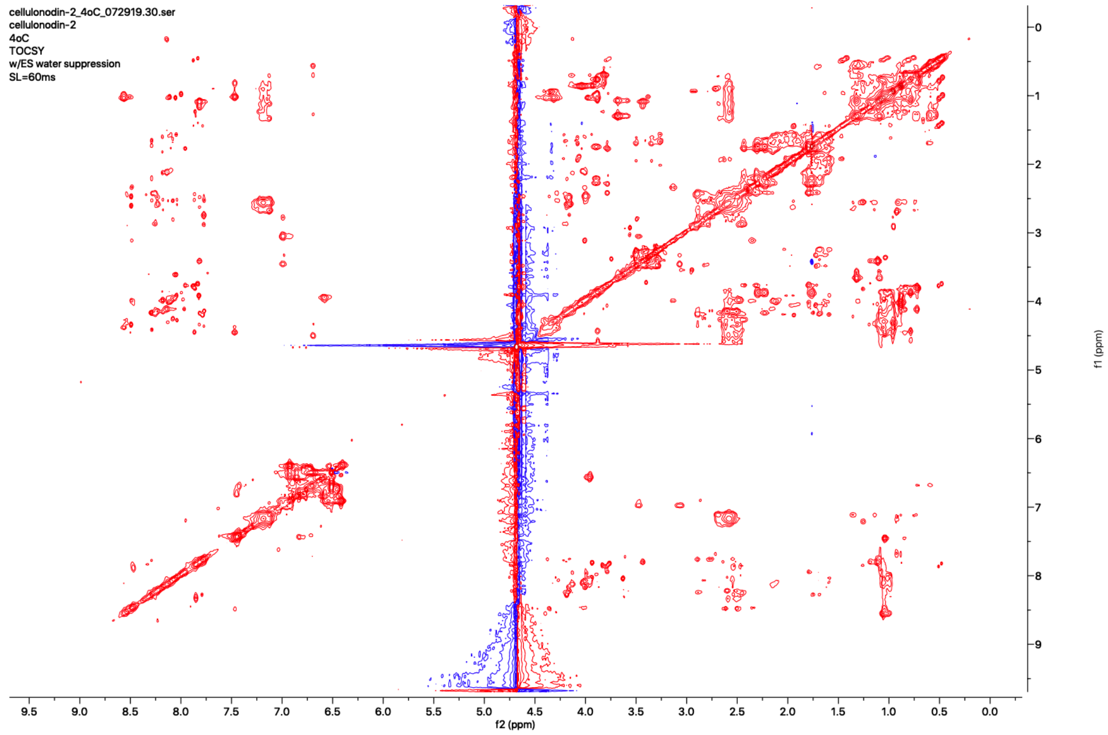
cellulonodin-2_4oC_072919.30.ser
cellulonodin-2
4oC
TOCSY
w/ES water suppression
SL=60ms
f2 (ppm)
f1 (ppm)

**c**

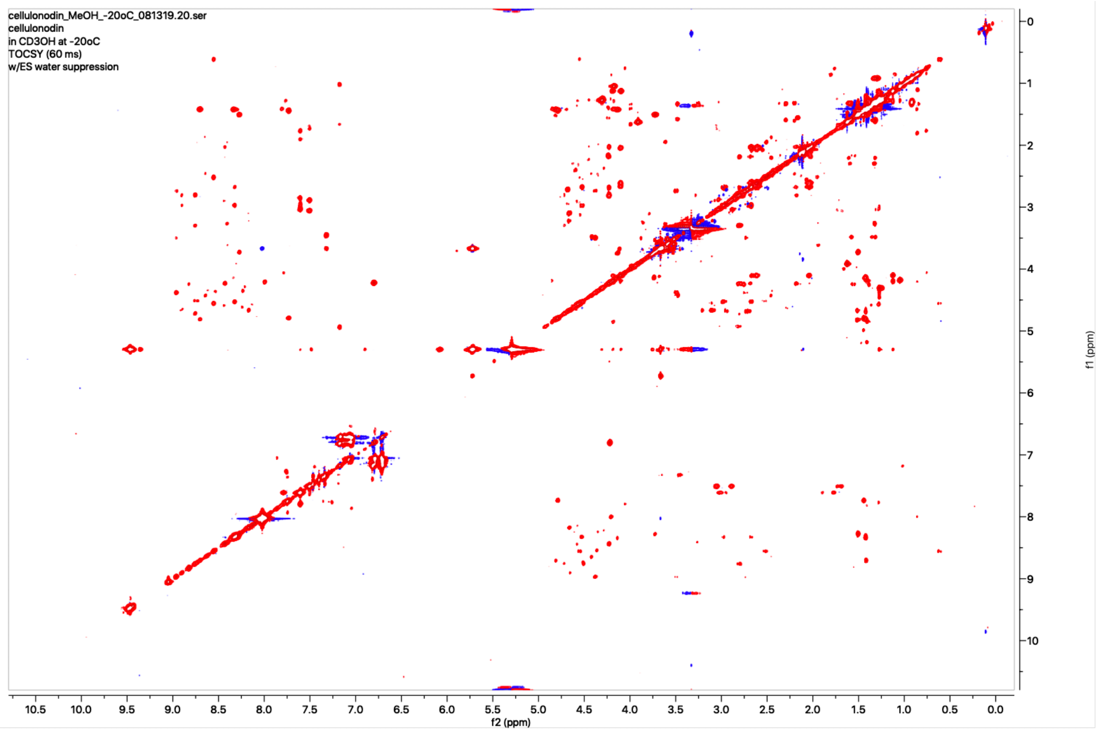

**Figure S10: TOCSY spectra of cellulonodin-2. a)** in 95:5 $H_2O$:$D_2O$ at 20 °C. **b)** in 95:5 $H_2O$:$D_2O$ at 4 °C. **c)** in $CD_3OH$ at -20 °C.

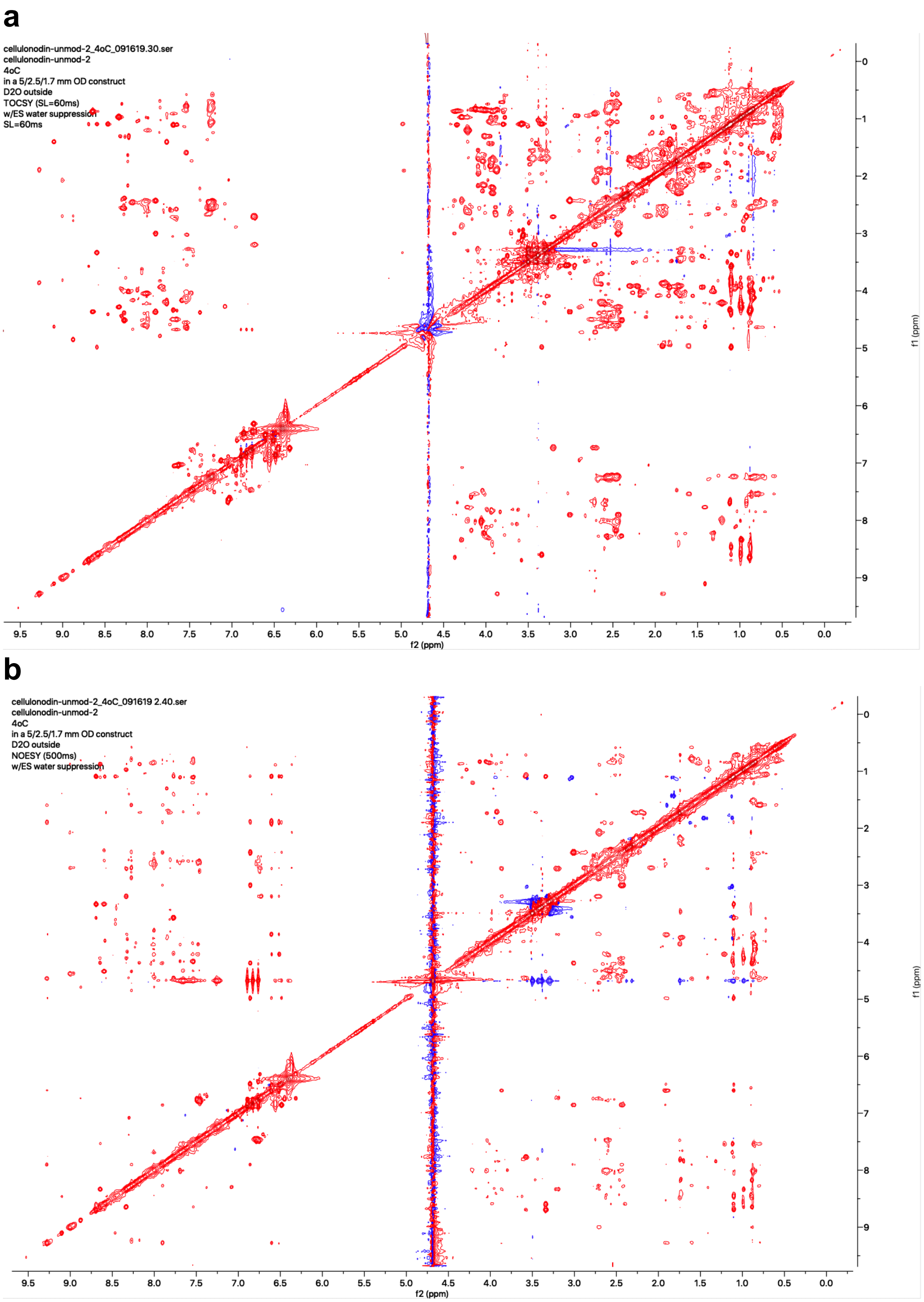

**Figure S11: 2D NMR spectra of lassoed only cellulonodin-2 at 4°C. a)** TOCSY. **b)** NOESY. Changes in the spectra during the acquisition suggested a dynamic structure, so structure calculations were not attempted.

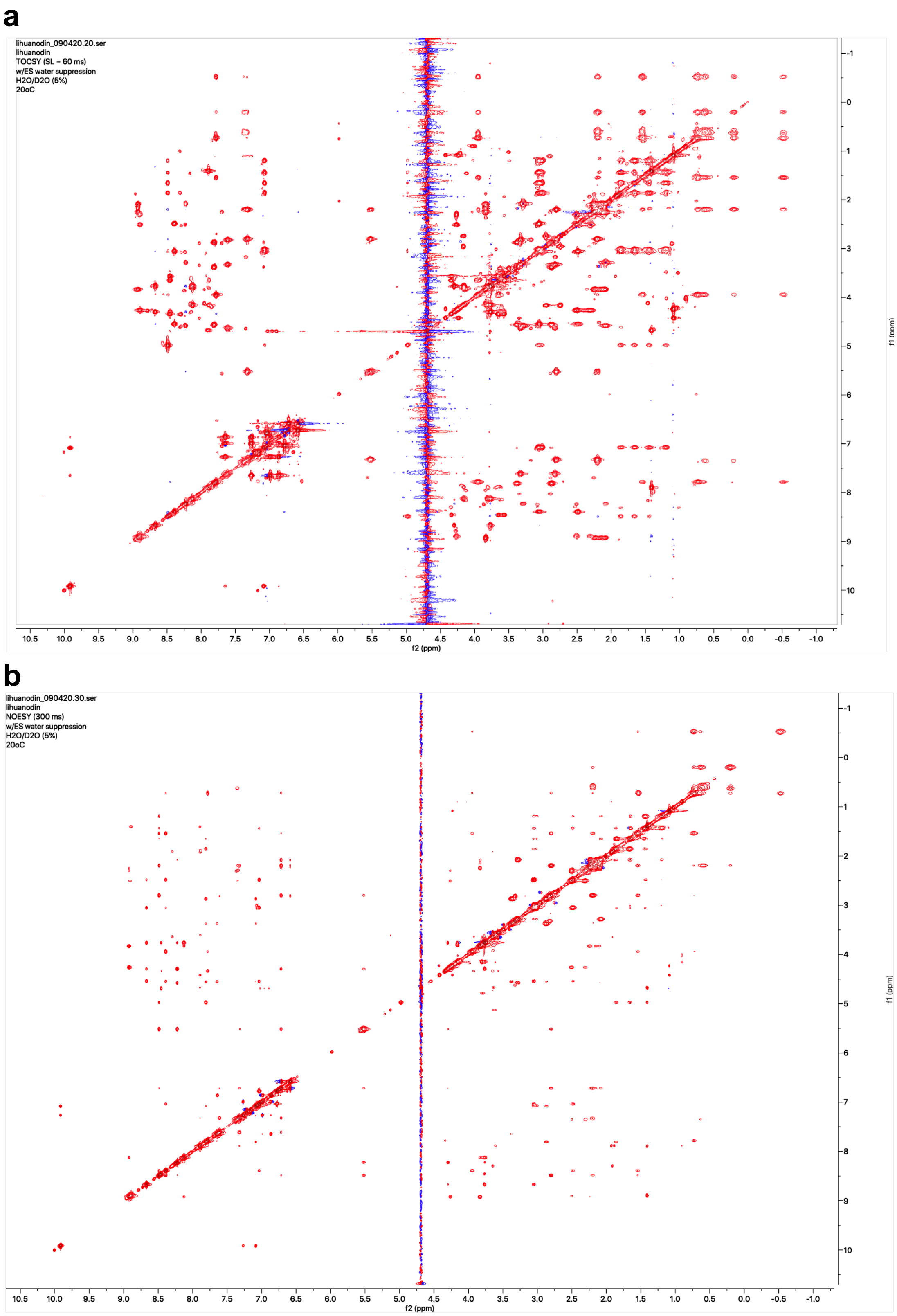

**Figure S12: 2D NMR of Lihuanodin. a)** TOCSY. **b)** NOESY. Peaks assignments are found in Table S2.

(a)

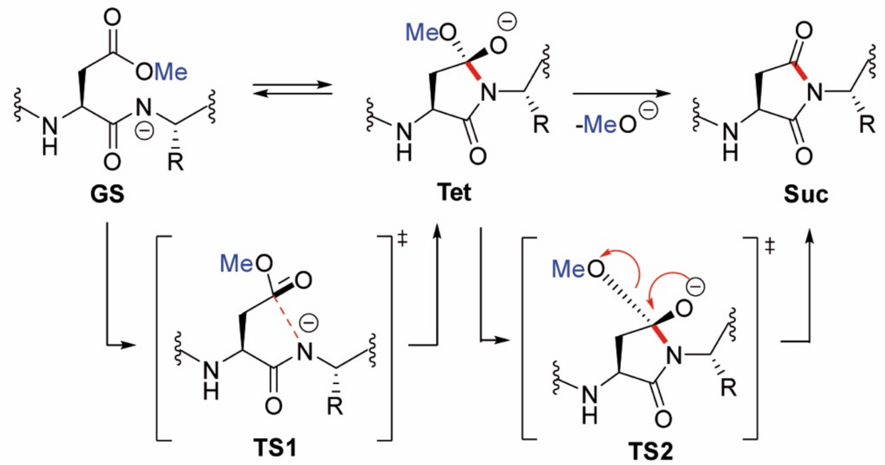

(b)

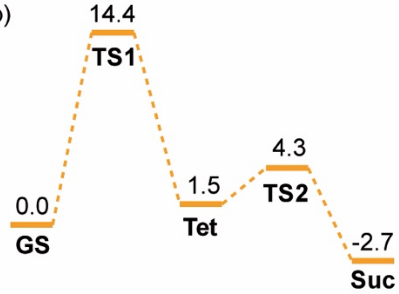

(c)

| Xxx | $\Delta E$ / kcal mol$^{-1}$ | $\Delta pK_a$ |
|---|---|---|
| Thr | 344.4 | 0 |
| Val | 350.5 | 2.7 |
| Ser | 339.9 | -0.2 |
| Ala | 348.7 | 3.4 |

**Figure S13. DFT Calculations on the Influence of Side Chain of *n*+1 Residue on Succinimide Formation. a)** Two-step mechanism of succinimide formation shown in its anionic form. **b)** Reaction coordinates with calculated ground and transition state energies in kcal/mol for the demethylation of AcAsp(OMe)ThrNHCH$_3$. **c)** Calculated gas-phase proton affinities (B3LYP-D3(BJ)/def2-TZVP) and $\Delta pK_a$ values (HF-3c/CPCM(water)) of the *n*+1 amide bond.

**a**

| | Aspartimidylated | Methylated | Lassoed | Linear core | Intensity of the major product |
|---|---|---|---|---|---|
| **WT** | 84.8% | 14.2% | 1.0% | 0.0% | $3.13 \times 10^7$ |
| **D5N** | | | Not observed | | |
| **D5T** | 67.2% | 17.1% | 15.6% | | $1.29 \times 10^7$ |
| **D5TD6N** | | | Not observed | | |
| **D6N** | 7.2% | / | 92.8% | 0.0% | $1.82 \times 10^6$ |
| **D6E** | / | 59.7% | 40.3% | 0.0% | $3.88 \times 10^6$ |
| **D6T** | 53.2% | 46.8% | / | 0.0% | $1.39 \times 10^6$ |
| **T7A** | 49.1% | 47.9% | 3.0% | 0.0% | $1.19 \times 10^7$ |
| **T7V** | / | 52.0% | 48.0% | 0.0% | $2.21 \times 10^7$ |
| **T7L** | / | 77.8% | 22.2% | 0.0% | $3.55 \times 10^7$ |
| **T7N** | 59.8% | 38.5% | 1.8% | 0.0% | $6.37 \times 10^6$ |
| **T7F** | 12.4% | 87.6% | / | 0.0% | $1.91 \times 10^7$ |
| **T7S** | 100.0% | 0.0% | 0.0% | 0.0% | $4.24 \times 10^6$ |
| **T7G** | 43.1% | 56.9% | 0.0% | 0.0% | $1.82 \times 10^5$ |
| **A8G** | 50.8% | 3.7% | 0.0% | 45.4% | $1.22 \times 10^6$ |
| **D9N** | | | Not observed | | |
| **N11D** | | | Not observed | | |
| **T12A** | 86.1% | 13.2% | 0.8% | 0.0% | $3.68 \times 10^7$ |
| **K14A** | | | Not observed | | |
| **K14Q** | | | Not observed | | |
| **K14F** | 4.4% | 92.0% | 3.6% | 0.0% | $2.37 \times 10^7$ |
| **Y15A** | | | Not observed | | |
| **Y15F** | 85.9% | 14.1% | / | 0.0% | $2.21 \times 10^7$ |
| **Y16F** | 95.5% | 1.9% | 2.6% | 0.0% | $1.22 \times 10^7$ |

**b**

| | Aspartimidylated | Methylated | Lassoed | Linear core | Intensity of the major product |
|---|---|---|---|---|---|
| **WT** | 100.0% | 0.0% | 0.0% | 0.0% | $3.37 \times 10^6$ |
| **D6N** | 0.0% | 0.0% | 100.0% | 0.0% | $2.31 \times 10^5$ |
| **T7A** | 100.0% | 0.0% | 0.0% | 0.0% | $1.10 \times 10^7$ |
| **T7V** | 75.4% | 24.6% | 0.0% | 0.0% | $8.65 \times 10^5$ |
| **T7L** | 100.0% | 0.0% | 0.0% | 0.0% | $3.96 \times 10^6$ |
| **A8G** | | | Not observed | | |
| **D9N** | | | Not observed | | |

**Figure S14: Species Detected in Cellulonodin-2 and Lihuanodin Variant Expressions, and Their Corresponding Percentages.** The intensity of the major product (AU from peak area in extracted ion chromatograms) for each mutant is also indicated. **a)** cellulonodin-2 variants. **b)** lihuanodin variants.

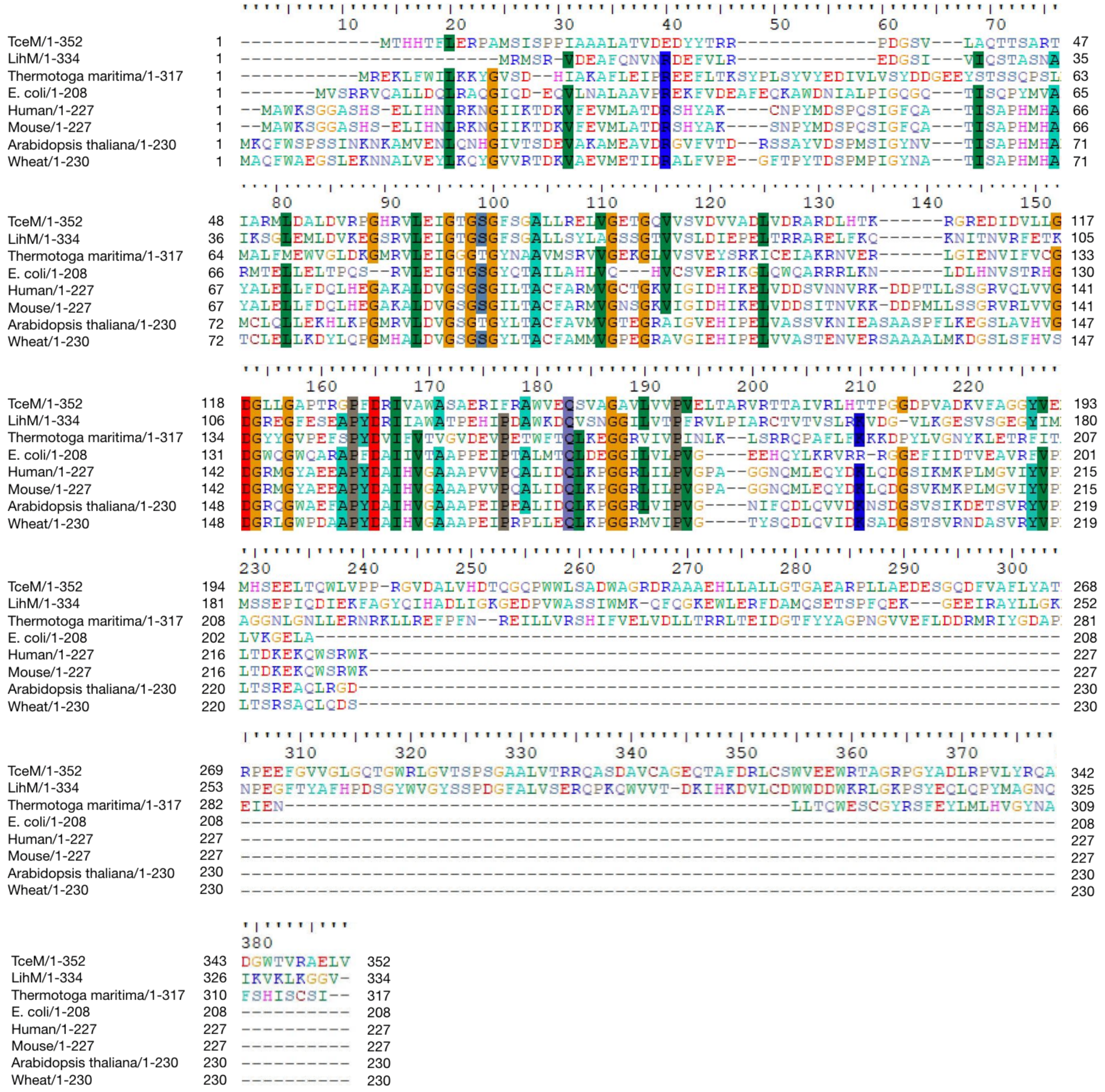

**Figure S15: Amino Acid Sequence Alignment of TceM and LihM with the Canonical Protein L-isoaspartate Methyltransferases (PIMTs) from Bacteria and Eukaryotes.**[18,19] Sequence alignment shows TceM and LihM consists of an N-terminal region that is homologous to canonical PIMTs, and a unique C-terminal segment. The one exception among the canonical PIMTs is *T. maritima*, which has a C-terminal extension of ~90 aa.

**a**

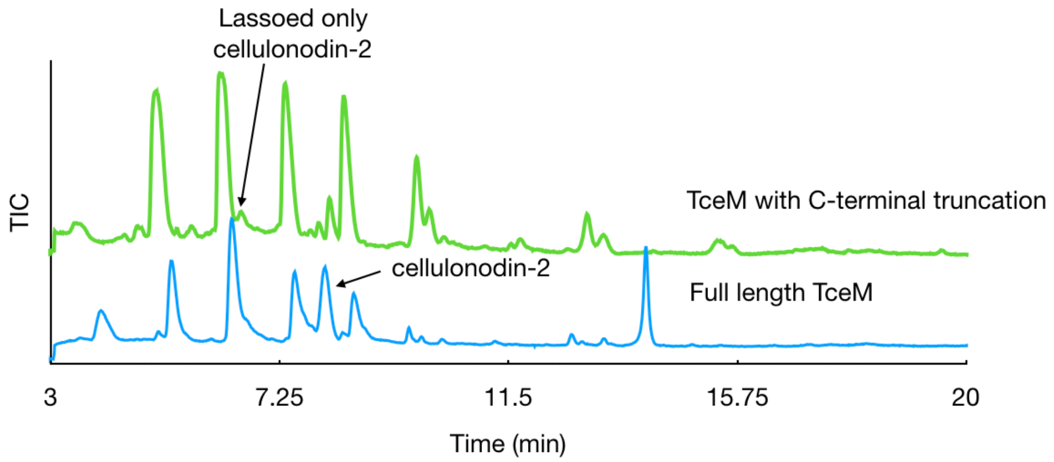

**b**

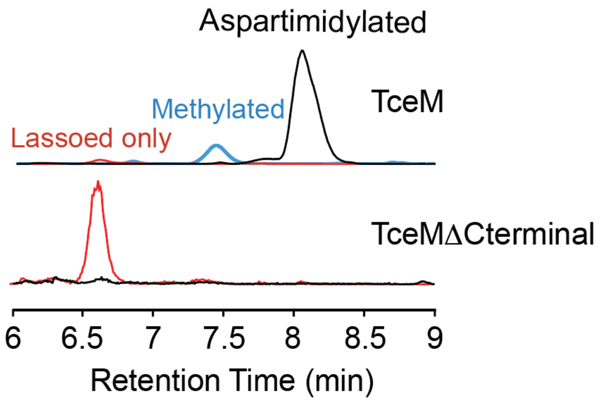

**c**

| | Aspartimidy -lated lasso | Methylated lasso | Lasso only | Intensity of the major product |
|---|---|---|---|---|
| **WT** | 84.8% | 14.2% | 1.0% | $3.1 \times 10^7$ |
| **C-terminal truncation** | 0.0% | 0% | 100% | $3.1 \times 10^5$ |

**Figure S16: Effect of TceM C-terminal Truncation *in vivo*. a)** TIC traces of cell extracts of cellulonodin-2 heterologous expressions with either full length TceM (blue) or TceM with 146 aa removed from its C-terminus (green). Removing the unique C-terminal segment of TceM abolishes its activity *in vivo.* No aspartimidylated lasso (cellulonodin-2) was observed. **b)** EIC of lassoed only (red), methylated (blue), and aspartimidylated (black) cellulonodin-2 using full-length TceM and TceMΔCterminal. **c)** product distributions and intensities from part b.

**a**

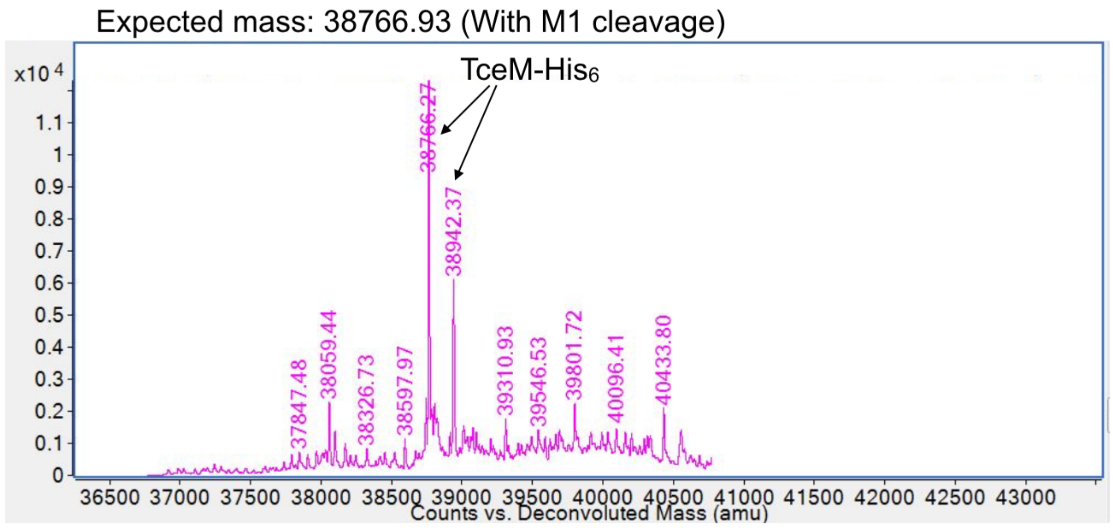

**b**

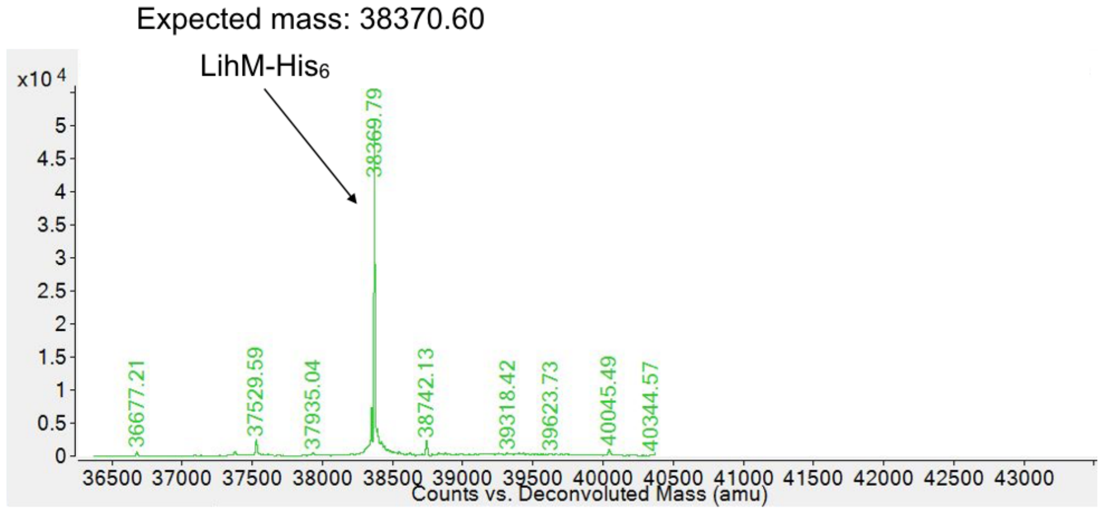

**Figure S17: Mass Spectra of TceM-His$_6$ and LihM-His$_6$. a)** TceM-His$_6$. The peak at 38766.27 Da corresponds to TceM-His$_6$ with first methionine cleaved. The species with a mass of 38942.37 corresponds to TceM_His$_6$ with the N-terminal methionine formylated. **b)** LihM-His$_6$. The indicated peak corresponds to full-length LihM-His$_6$ with the N-terminal methionine intact.

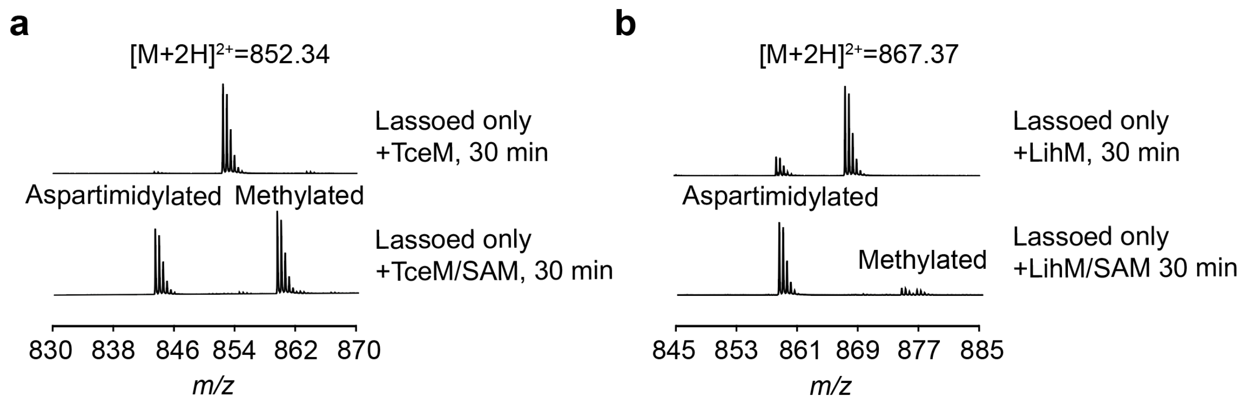

**Figure S18: Comparison between *In vitro* Reactions with and without SAM.** Reactions include 100 mM lassoed only substrate, 1 mM TceM or LihM, 1 mM SAM (if present) in 50 mM Tris-HCl at pH 7.4. Reactions carried out at 37 °C. **a)** cellulonodin-2. **b)** lihuanodin. Little or no aspartimidylation is observed without SAM, as expected.

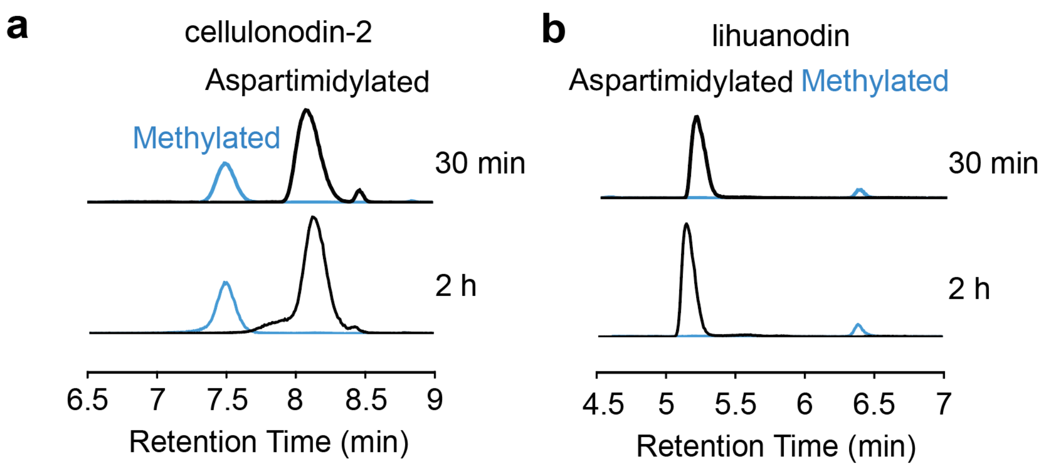

**Figure S19: Comparison between *In vitro* Reactions for 30 mins and 2 hours for Cellulonodin-2 and Lihuanodin. a)** Extracted ion chromatograms (EIC) for cellulonodin-2 species. **b)** EICs for lihuanodin species. The peaks for aspartimidylated and methylated species are labelled black and blue respectively. Extending the reaction beyond 30 min has little effect on the product distribution.

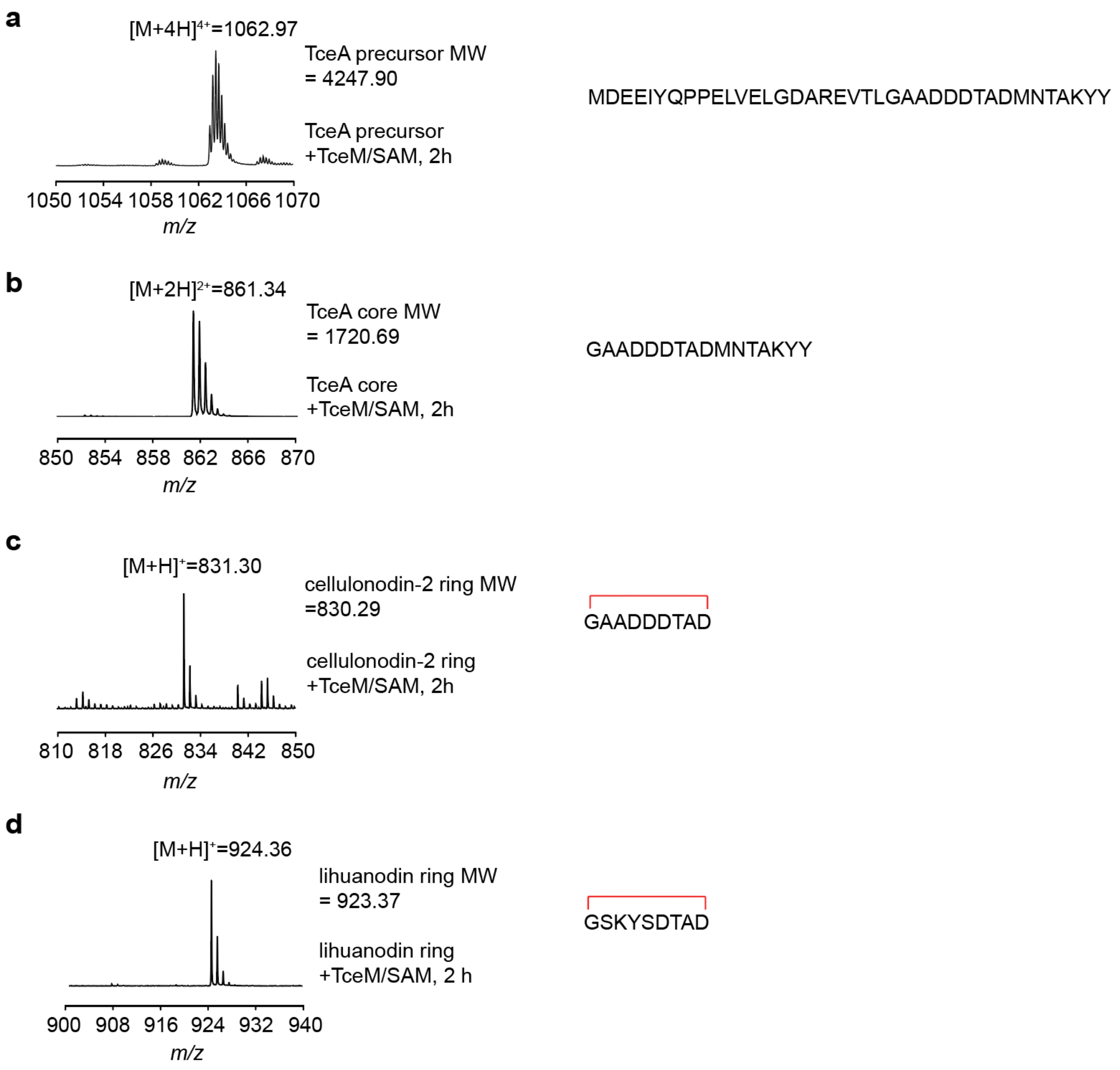

**Figure S20: Identification of Substrates for TceM and LihM. a)** *In vitro* reaction of TceA linear precursor with TceM for 2 h. **b)** *In vitro* reaction of TceA linear core with TceM for 2 h. **c)** *In vitro* reaction of cellulonodin-2 isopeptide bonded ring with TceM for 2 h. **d)** *In vitro* reaction of lihuanodin isopeptide bonded ring with LihM for 2 h. No reaction observed in all of these cases suggesting that the entire lasso structure is required for recognition by TceM and LihM.

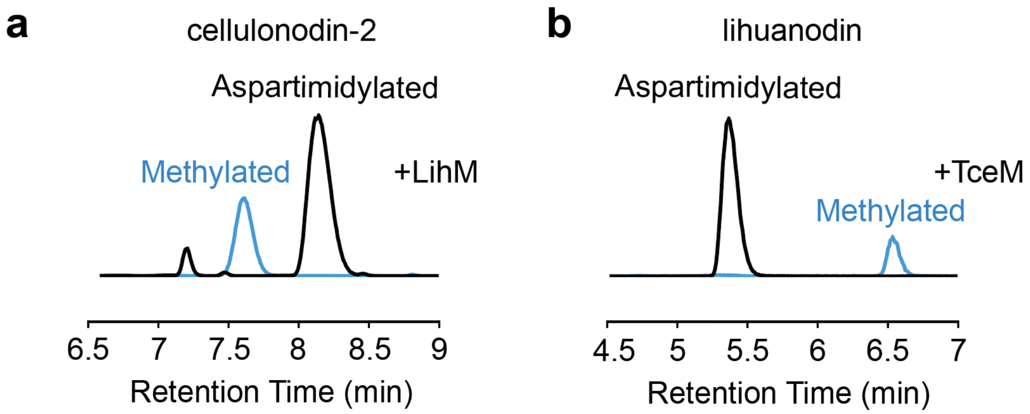

**Figure S21: *In vitro* Cross Reaction of TceM/LihM with Lassoed Only Lihuanodin/ Cellulonodin-2. a)** EICs of lassoed only cellulonodin-2 reacted with LihM. **b)** EICs of lassoed only lihuanodin reacted with TceM. The aspartimidylated and methylated peaks were labelled black and blue respectively.

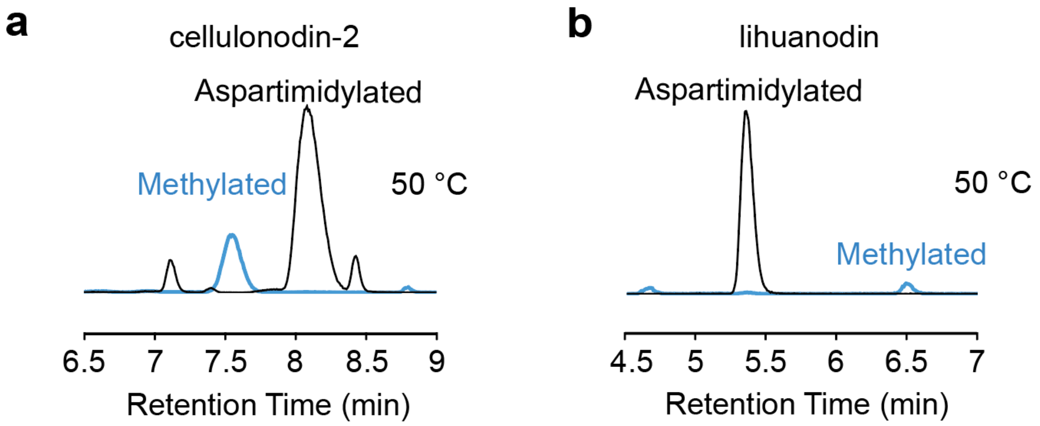

**Figure S22:** ***In vitro*** **Reactions at 50 °C. a)** EICs of cellulonodin-2. **b)** EICs of lihuanodin. The aspartimidylated and methylated peaks were labelled black and blue respectively. The product compositions for both reactions were similar to those in *in vitro* reactions at 37 °C (Figure 8 in the main text).

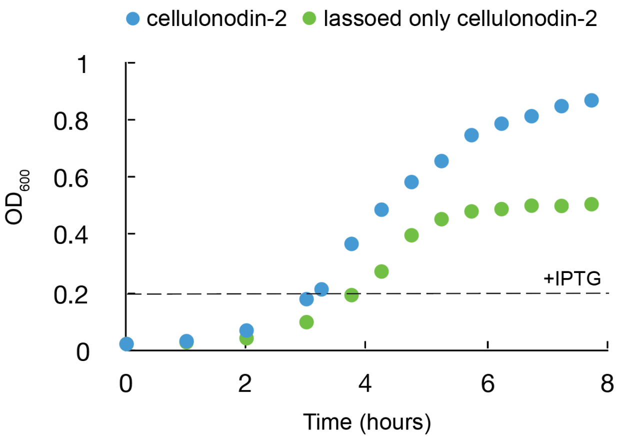

**Figure S23: Growth curves of *E. coli* BL21 expressing cellulonodin-2 (blue) and lassoed only cellulonodin-2 (green).** The expression of peptides was induced using 1 mM IPTG when the $OD_{600}$ of the culture reached 0.2. Lassoed only cellulonodin-2 caused a lower final culture OD compared to aspartimidylated cellulonodin-2.

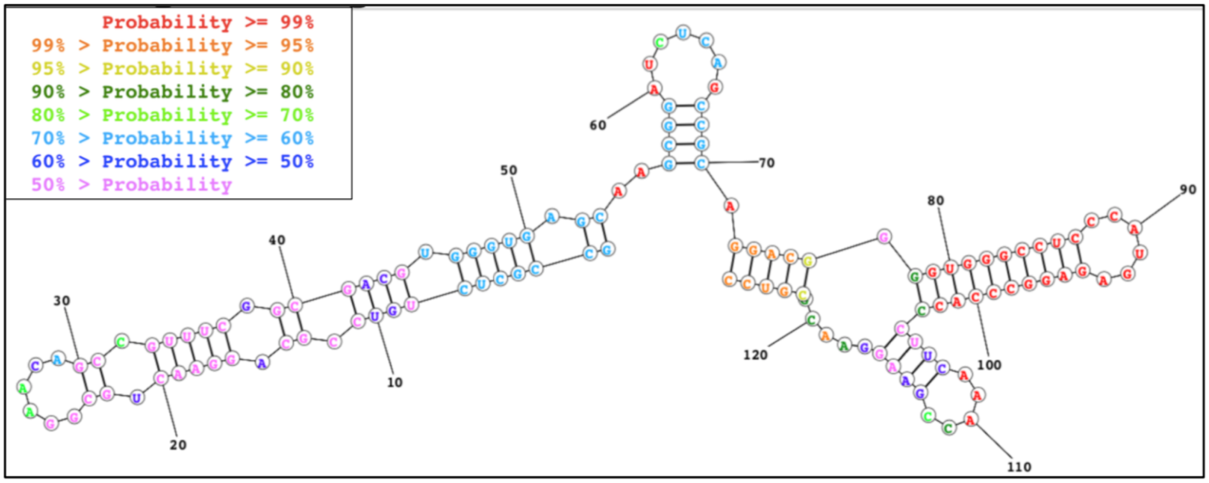

**Figure S24: RNA Structure of the Intergenic Region of the Cellulonodin-2 Gene Cluster Predicted by *RNAStructure*.** The predicted structure with the lowest free energy structure of the intergenic region between *TceA* and *TceC* exhibits multiple hairpin regions predicted at high probability. The inset color legend describes the percent probability of each nucleotide making up the predicted structure. This 126 base-pair intergenic region is indicated in **Figure 1**.

**a**

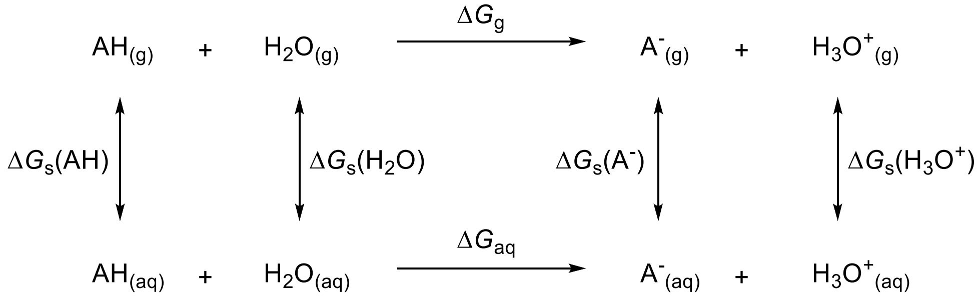

**b**

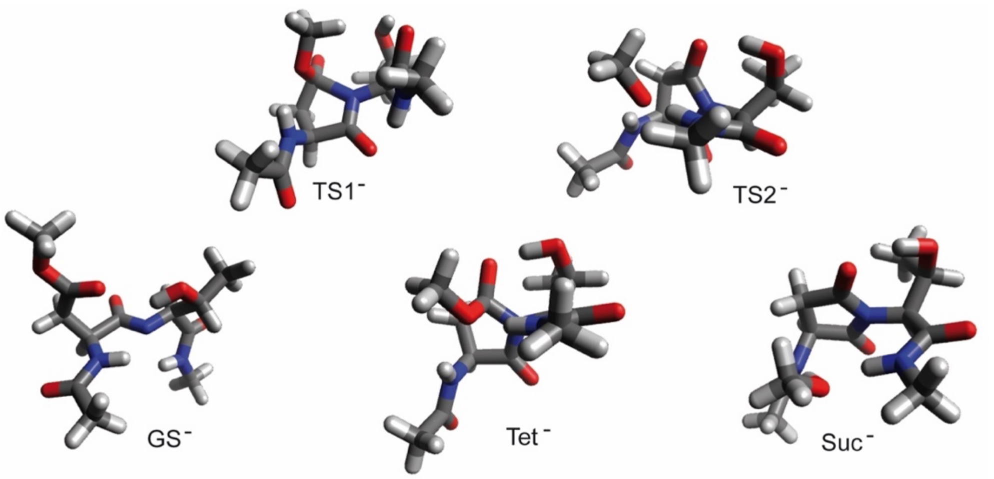

**Figure S25: Further Details About Asp-Xxx DFT Calculations. a**) Proton exchange thermodynamic cycle for calculations of $\Delta pK_a$ values. **b)** Optimized minimum and transition state structures of model dipeptide AcAsp($OCH_3$)Thr$NHCH_3$ after deprotonation of the central amide bond.

### Supplementary Tables

#### Table S1. Primers and constructs used in this study.

| Primer Name | Primer Sequence | Primer Description | Plasmid construct | Plasmid Description |
|---|---|---|---|---|
| JK131 | TATTCTAGATTGCTGCGGAAGGAACATATGGACG AGGAGATC | Forward primer for *T.cellulosilytica* cluster 2 | pASK75 | pJDK87/pMB4,p MB7-19 |
| JK132 | TATAAGCTTACGGTCAGACCAACTCCGCGCGCA | Reverse primer for *T.cellulosilytica* cluster 2 (with TceM) | pASK75 | pJDK87 |
| JK159 | TGACAGCTGTCAAGCCGGCCT | *Pvu*II forward for T. cell B2-2 change to ATG | pASK75 | pJDK92 |
| JK160 | GGACAGCCAAaTGAGCATGCA | *tce*B2-2 ATG forward | pASK75 | pJDK92 |
| JK161 | TGCATGCTCAtTTGGCTGTCC | *tce*B2-2 ATG reverse | pASK75 | pJDK92 |
| JK162 | AGGGAATTCTCCAGACAGGC | *Eco*RI reverse for T. cell B2-2 change to ATG | pASK75 | pJDK92 |
| JK169 | ATCTCGAGAGAGGAGAAATTAACTATGGCGATAC GGAAGTTCTT | Remove *tce* hairpin forward | pASK75 | pMB4 |
| JK170 | GTTAATTTCTCCTCTCTCGAGATACTAGTAGTACT TGGCCGTGT | Remove *tce* hairpin reverse | pASK75 | pMB4 |
| JK171 | GGTGCTGAGCTGCGAGCAGT | *tceC Blp*I reverse | pASK75 | pMB4 |
| JK208 | CGTccATGgCCCACCACACCTTCTTGGAA | *T. cellulosilytica* methyltransferase for MCS1-Forward | pRSF-Duet | pJDK127, pLC40-41 |
| JK209 | ataaagcttTCAgtgatggtgatggtgatg | *T. cellulosilytica* methyltransferase for MCS1-Reverse | pRSF-Duet | pJDK127, pLC40-41 |
| MCB1 | ACAGAATTCATTAAAGAGGAGAAATTAACTATGG ACGAGGAGATCTATCAGCCCC | Forward *tceA* (With *Eco*RI+RBS) | pQE80 | pMB2 |
| MCB2 | TTTAAGCTTCTAGTAGTACTTGGCCGTGTTCATG | Reverse *tceA* | pQE80 | pMB2 |
| MCB6 | ATGGCGATACGGAAGTTCTTCATCG | Forward *tceC* | pQE80 | pMB3 |
| MCB4 | TTTGCCCATGGTCAGACCAACTCCGCGCGCA | Reverse *tceM* | pQE80 | pMB3 |
| MCB7 | CTCTCTCGAGATACTAGTAGTACTTGGCCGTGTT CATGTCGGCGGTGTTGTCGTCCGCCG | Reverse TceA D6N | pASK75 | pMB7 |
| MCB8 | CTCTCTCGAGATACTAGTAGTACTTGGCCGTGTT CATGTTGGCGGTGTCGTCGTCCGCCG | Reverse TceA D9N | pASK75 | pMB11 |
| MCB9 | CTCTCTCGAGATACTAGTAGTACTTGGCCGTGTT CATGTCGGCGGCGTCGTCGTCCGCCG | Reverse TceA T7A | pASK75 | pMB15 |
| MCB10 | CTCTCTCGAGATACTAGTAGTACTTGGCCGTGTT CATGTCGGCGGTGGTGTCGTCCGCCG | Reverse TceA D6T | pASK75 | pMB8 |
| MCB11 | CTCTCTCGAGATACTAGTAGTACTTGGCCGTGTC CATGTCGGCGGTGTCGTCGTCCGCCG | Reverse TceA N11D | pASK75 | pMB12 |
| MCB12 | CTCTCTCGAGATACTAGTAGTACTTGGCCGTGTT CATGTCGGCCACGTCGTCGTCCGCCG | Reverse TceA T7V | pASK75 | pMB16 |
| MCB13 | CTCTCTCGAGATACTAGTAGTACTTGGCCGTGTT CATGTCGGCCAGGTCGTCGTCCGCCG | Reverse TceA T7L | pASK75 | pMB17 |
| MCB14 | CTCTCTCGAGATACTAGTAGTACTTGGCCGTGTT CATGTCGGCGTTGTCGTCGTCCGCCG | Reverse TceA T7N | pASK75 | pMB18 |
| MCB15 | CTCTCTCGAGATACTAGTAGTACTTGGCCGTGTT CATGTCGGCGAAGTCGTCGTCCGCCG | Reverse TceA T7F | pASK75 | pMB19 |
| MCB18 | CTCTCTCGAGATACTAGTAGTACTTGGCCGTGTT CATGTCGGCGGTGTCGGTGTCCGCCG | Reverse TceA D5T | pASK75 | pMB9 |

| | | | | |
|---|---|---|---|---|
| MCB19 | CTCTCTCGAGATACTAGTAGTACTTGGCCGTGTTCATGTCGGCGGTGTTGGTGTCCGCCG | Reverse TceA D5TD6N | pASK75 | pMB10 |
| MCB20 | CTCTCTCGAGATACTAGTAGTACGCGGCCGTGTTCATGTCGGCGGTGTCGTCGTCCGCCG | Reverse TceA K14A | pASK75 | pMB13 |
| MCB21 | CTCTCTCGAGATACTAGTAGGCCTTGGCCGTGTTCATGTCGGCGGTGTCGTCGTCCGCCG | Reverse TceA Y15A | pASK75 | pMB14 |
| MO1 | TGCATCTAGATTGCTGCGGAAGGA | TceA *Xba*I Forward | pASK75 | pMO1-6 |
| MO2 | GTCAGCTGAGCTGCGAGCAG | TceA *Blp*I Reverse | pASK75 | pMO1-6 |
| MO3 | ACGACGAAACCGCCGA | Forward TceA D6E | pASK75 | pMO1 |
| MO4 | TCGGCGGTTTCGTCGT | Reverse TceA D6E | pASK75 | pMO1 |
| MO5 | CACGGCCAAGTTCTACTAGTATCTC | Forward TceA Y15F | pASK75 | pMO2 |
| MO6 | GAGATACTAGTAGAACTTGGCCGTG | Reverse TceA Y15F | pASK75 | pMO2 |
| MO7 | CGACATGAACgCGGCCAA | Forward TceA T12A | pASK75 | pMO3 |
| MO8 | TTGGCCGcGTTCATGTCG | Reverse TceA T12A | pASK75 | pMO3 |
| MO9 | TGAACACGGCCcAGTACTACTAG | Forward TceA K14Q | pASK75 | pMO4 |
| MO10 | CTAGTAGTACTgGGCCGTGTTCA | Reverse TceA K14Q | pASK75 | pMO4 |
| MO11 | ACATGAACACGGCCttcTACTACTAGTA | Forward TceA K14F | pASK75 | pMO5 |
| MO12 | TACTAGTAGTAgaaGGCCGTGTTCATGT | Reverse TceA K14F | pASK75 | pMO5 |
| MO13 | CACGGCCAAGTACttCTAGTATCTC | Forward TceA Y16F | pASK75 | pMO6 |
| MO14 | GAGATACTAGaaGTACTTGGCCGTG | Reverse TceA Y16F | pASK75 | pMO6 |
| MO15 | GCGATCTAGAATGAGAATGAGTCGAGTTGATGAAG | *L. thermophila* cluster *Xba*I Forward | pASK75 | pMO7 |
| MO16 | TCGCGTCGACCTATACTACTATAATTTTTTTATATGCCTTAATTTTCTCTGCTTCT | *L. thermophila* cluster *Sal*I Reverse | pASK75 | pMO7 |
| MO17 | GTGCGAATTCATTAAAGAGGAGAAATTAACTATGGAGTACCAAGTGCCC | LihA *Eco*RI RBS Forward | pQE80 | pMO8 |
| MO18 | TCGCAAGCTTTTACCAACGGTAGGAGCTTTC | LihA *Hind*III Reverse | pQE80 | pMO8 |
| LC1 | TCCAAGCTAGCcatcaattaagaaaaa | mcjBCD promoter *Nhe*I Forward | pQE80 | pMO9 |
| MO19 | ggagtgtgcaaaATGTTTCTTGGAATTTATGAGGGTTATTCT | LihC internal forward | pQE80 | pMO9 |
| MO20 | CCAAGAAACATtttgcacactccctcctg | LihC internal reverse | pQE80 | pMO9 |
| MO21 | TCGCCCATGGCTATACTACTATAATTTTTTTATATGCCTTAATTTTCTCTGCT | LihB2 *Nco*I Reverse | pQE80 | pMO9 |
| LC2 | ccaaGCTAGCCATCAATTAAGAAAA | Lassoed only *Nhe*I forward | pQE80 | pLC13 |
| LC3 | AGACCCATGGTCATACACGAAGCAGCAGTA | Lassoed only NcoI Reverse | pQE80 | pLC13 |
| LC4 | CTTCACCTCGAGAAATCATAAAAAATTTATTTGC | Tce/Lih mutant pQE80 *Xho*I forward | pQE80 | pLC14-18, 32-35, 37-39 |
| LC5 | GCTAGCTTGGATTCTCACCAATAAA | Tce/Lih mutant pQE80 *Nhe*I Reverse | pQE80 | pLC14-18, 32-35, 37-39 |
| LC6 | CGGCGGACACCGACA | TceA D5T forward pQE80 | pQE80 | pLC14 |
| LC7 | TGTCGGTGTCCGCCG | TceA D5T reverse pQE80 | pQE80 | pLC14 |
| LC8 | GACGACACCACCGCCG | TceA D6T forward pQE80 | pQE80 | pLC15 |
| LC9 | CGGCGGTGGTGTCGTC | TceA D6T reverse pQE80 | pQE80 | pLC15 |
| LC10 | GACGACAACGCCGAC | TceA D7N forward pQE80 | pQE80 | pLC16 |
| LC11 | GTCGGCGTTGTCGTC | TceA D7N reverse pQE80 | pQE80 | pLC16 |

| | | | | |
|---|---|---|---|---|
| LC12 | TTTAAGCTTCTAGTAGTAAAAGGCCGTG | TceA K14F reverse pQE80 | pQE80 | pLC17 |
| LC13 | TTTAAGCTTCTAGTAGTACTGGGCCG | TceA K14Q reverse pQE80 | pQE80 | pLC18 |
| LC14 | GCTGGTGCCGCCCTAACCATGGAGAT | TceM truncation NcoI reverse | pQE80 | pLC26 |
| LC15 | GACGACTCAGCCGAC | TceA T7S forward pQE80 | pQE80 | pLC27 |
| LC16 | GTCGGCTGAGTCGTC | TceA T7S reverse pQE80 | pQE80 | pLC27 |
| LC17 | CGACACCGgCGACATG | TceA A8G forward Pqe80 | pQE80 | pLC28 |
| LC18 | CATGTCGcCGGTGTCG | TceA A8G reverse pQE80 | pQE80 | pLC28 |
| LC19 | CACAGAATTCATTAAAGAGGAGAAATTAACTATG AGAATGAGTCGAGTTGATGAAGCTTT | LihM His6 *Eco*RI forward | pQE80 | pLC29 |
| LC20 | TGCAGGTCGACTTAGTGATGGTGATGGTGATGT ACACCTCCTTTCAATTTCACTTTGA | LihM His6 *Sal*I Reverse | pQE80 | pLC29 |
| LC21 | AGTCGGTCTCACATGAGAATGAGTCGAGTTGAT GAAG | LihM *Bsa*I forward | pRSF Duet | pLC30, 42-43 |
| LC22 | GCTCGTCGACTTATACACCTCCTTTCAATTTCAC TTTGATTT | LihM *Sal*I Reverse | pRSF Duet | pLC30, 42-43 |
| LC23 | GTTCCAAATATAGCAATACAGCAGATG | LihA D6N forward | pQE80 | pLC32 |
| LC24 | CATCTGCTGTATTGCTATATTTGGAAC | LihA D6N reverse | pQE80 | pLC32 |
| LC25 | CCAAATATAGCGATGCAGCAGATGAA | LihA T7A forward | pQE80 | pLC33 |
| LC26 | TTCATCTGCTGCATCGCTATATTTGG | LihA T7A reverse | pQE80 | pLC33 |
| LC27 | CAAATATAGCGATACAGGAGATGAAAGCT | LihA A8G forward | pQE80 | pLC34 |
| LC28 | AGCTTTCATCTCCTGTATCGCTATATTTG | LihA A8G reverse | pQE80 | pLC34 |
| LC29 | CGATACAGCAAATGAAAGCTCC | LihA D9N forward | pQE80 | pLC35 |
| LC30 | GGAGCTTTCATTTGCTGTATCG | LihA D9N reverse | pQE80 | pLC35 |
| LC31 | GATGATGGCGCGGAT | TceA T7G forward | pQE80 | pLC37 |
| LC32 | GATGATGGCGCGGAT | TceA T7G forward | pQE80 | pLC37 |
| LC33 | GTGGGCGCGGTGGTG | LihA T7L forward | pQE80 | pLC38 |
| LC34 | CCAAATATAGCGATTTAGCAGATGAA | LihA T7L reverse | pQE80 | pLC38 |
| LC35 | TTCATCTGCTAAATCGCTATATTTGG | LihA T7V forward | pQE80 | pLC39 |
| LC36 | CCAAATATAGCGATGTAGCAGATGAA | LihA T7V reverse | pQE80 | pLC39 |
| LC37 | TTCATCTGCTACATCGCTATATTTGG | TceM W323A forward | pRSF Duet | pLC40 |
| LC38 | GGAGGAAGCGAGGACAGC | TceM W323A reverse | pRSF Duet | pLC40 |
| LC39 | GGTCGGGCAGGCTAC | TceM P329A forward | pRSF Duet | pLC41 |
| LC40 | TGTCCTCTTCCTCCacc | TceM P329A reverse | pRSF Duet | pLC41 |
| LC41 | GGATGACGCGAAACGGCTT | LihM W306A forward | pRSF Duet | pLC42 |
| LC42 | CCGTTTCgcGTCATCCCac | LihM W306A reverse | pRSF Duet | pLC42 |
| LC43 | GGCTTGGAAAggCAAGTTATGA | LihM P312A forward | pRSF Duet | pLC43 |
| LC44 | TCATAACTTGCCTTTCCAAGCC | LihM P312A reverse | pRSF Duet | pLC43 |

#### Table S2. Chemical shift assignments for lihuanodin

| Residue | Hydrogen | Chemical shift δ (ppm) | Residue | Hydrogen | Chemical shift δ (ppm) |
|---|---|---|---|---|---|
| GLY-1 | H | 7.347 | GLU-10 | H | 8.924 |
| GLY-1 | QA | 4.638 | GLU-10 | HA | 3.834 |
| SER-2 | H | 8.459 | GLU-10 | QB | 2.238 |
| SER-2 | HA | 4.339 | GLU-10 | HG2 | 2.150 |
| SER-2 | HB2 | 3.652 | GLU-10 | HG3 | 2.085 |
| SER-2 | HB3 | 3.564 | SER-11 | H | 8.127 |
| LYS-3 | H | 7.783 | SER-11 | HA | 4.151 |
| LYS-3 | HA | 3.946 | SER-11 | HB2 | 3.811 |
| LYS-3 | QB | 1.545 | SER-11 | HB3 | 3.776 |
| LYS-3 | HD2 | 0.723 | SER-12 | H | 7.613 |
| LYS-3 | HD3 | 0.606 | SER-12 | HA | 4.627 |
| LYS-3 | QE | 2.199 | SER-12 | HB2 | 3.326 |
| LYS-3 | HG2 | 0.201 | SER-12 | HB3 | 2.819 |
| LYS-3 | HG3 | -0.521 | TYR-13 | H | 7.324 |
| TYR-4 | H | 8.392 | TYR-13 | HA | 5.519 |
| TYR-4 | HA | 4.545 | TYR-13 | HB2 | 2.801 |
| TYR-4 | HB2 | 3.054 | TYR-13 | HB3 | 2.202 |
| TYR-4 | HB3 | 2.484 | TYR-13 | QD | 6.717 |
| TYR-4 | QD | 6.781 | TYR-13 | QE | 6.588 |
| TYR-4 | QE | 7.035 | ARG-14 | H | 8.488 |
| SER-5 | H | 8.671 | ARG-14 | HA | 4.979 |
| SER-5 | HA | 4.298 | ARG-14 | HB2 | 1.657 |
| SER-5 | QB | 3.764 | ARG-14 | HB3 | 1.856 |
| ASP-6 | H | 8.227 | ARG-14 | HG2 | 1.199 |
| ASP-6 | HA | 4.580 | ARG-14 | HG3 | 1.439 |
| ASP-6 | HB2 | 3.288 | ARG-14 | HD2 | 3.007 |
| ASP-6 | HB3 | 2.079 | ARG-14 | HD3 | 3.065 |
| THR-7 | HA | 4.424 | ARG-14 | HE | 7.071 |
| THR-7 | HB | 4.233 | TRP-15 | H | 7.811 |
| THR-7 | QG | 1.087 | TRP-15 | HA | 4.556 |
| ALA-8 | H | 7.896 | TRP-15 | HB2 | 2.872 |
| ALA-8 | HA | 4.674 | TRP-15 | HB3 | 3.376 |
| ALA-8 | QB | 1.410 | TRP-15 | HD | 7.086 |
| ASP-9 | H | 8.896 | TRP-15 | HE1 | 9.918 |
| ASP-9 | HA | 4.263 | TRP-15 | HE3 | 7.644 |
| ASP-9 | HB2 | 2.508 | TRP-15 | HH2 | 6.992 |
| ASP-9 | HB3 | 2.302 | TRP-15 | HZ2 | 7.268 |
| | | | TRP-15 | HZ3 | 6.863 |

#### Table S3. List of Through Space NOE Correlations between Ring and Lock Residues

| Position | Residue | Hydrogen | Position | Residue | Hydrogen |
|---|---|---|---|---|---|
| 13 | Tyr | HA | 5 | Ser | H |
| 13 | Tyr | HA | 6 | Asp | H |
| 13 | Tyr | HB3 | 4 | Tyr | H |
| 13 | Tyr | HB3 | 6 | Asp | H |
| 13 | Tyr | QD | 3 | Lys | QB |
| 13 | Tyr | QD | 3 | Lys | HD2 |
| 13 | Tyr | QD | 3 | Lys | HG2 |
| 13 | Tyr | QD | 4 | Tyr | H |
| 13 | Tyr | QD | 5 | Ser | H |
| 13 | Tyr | QD | 5 | Ser | HA |
| 13 | Tyr | QD | 6 | Asp | H |
| 13 | Tyr | QE | 5 | Ser | HA |
| 14 | Arg | H | 3 | Lys | HA |
| 14 | Arg | H | 3 | Lys | QB |
| 14 | Arg | H | 3 | Lys | HD2 |
| 14 | Arg | H | 5 | Ser | HA |
| 14 | Arg | HG2 | 4 | Tyr | H |
| 15 | Trp | H | 8 | Ala | QB |

**Table S4.** Calculated single point energies (SPE) and gas-phase proton affinities (**Δ*E***) of model compounds AcAsp($OCH_3$)XxxNHC$H_3$

| Xxx | **Ground state SPE**[a] / $E_h$ | **anionic ground state SPE**[a] / $E_h$ | **Δ*E*** = $E_{anionic}$ - $E_{neutral}$ / kcal·mol$^{-1}$ |
|---|---|---|---|
| Thr | -1085.5047 | -1084.9559 | 344.4 |
| Val | -1049.5875 | -1049.0290 | 350.5 |
| Ser | -1046.1959 | -1045.6542 | 339.9 |
| Ala | -970.9762 | -970.4206 | 348.7 |

[a] single point energies were performed on optimized structures at the B3LYP-D3(BJ)/def2-TZVP level

**Table S5.** Calculated thermodynamic parameters and p$K_a$ values of model dipeptides AcAsp($OCH_3$)XxxNHC$H_3$ and their corresponding anions

| Xxx | electronic energy[a] / $E_h$ | $E^{ZEP}$ + $E^{vib}$ + $E^{rot}$ + $E^{trans}$ / $E_h$ | $H$ / $E_h$ | -T·$S$ / $E_h$ | $G$[b] / $E_h$ | $\Delta G_s$[c] / kcal·mol$^{-1}$ | $\Delta G_{aq}$ / kcal·mol$^{-1}$ | p$K_a$[d] |
|---|---|---|---|---|---|---|---|---|
| Thr | -1072.2557 | 0.4124 | -1071.8423 | -0.0730 | -1071.9153 | -21.8 | 67.4 | 47.7 |
| Val | -1036.6760 | 0.4435 | -1036.2315 | -0.0745 | -1036.3061 | -17.0 | 71.0 | 50.4 |
| Ser | -1033.4657 | 0.3813 | -1033.0835 | -0.0737 | -1033.1572 | -23.2 | 67.2 | 47.5 |
| Ala | -959.0970 | 0.3745 | -958.7216 | -0.0702 | -958.7917 | -18.0 | 72.0 | 51.1 |
| Thr$^-$ | -1071.6872 | 0.3995 | -1071.2867 | -0.0731 | -1071.3598 | -59.8 | / | / |
| Val$^-$ | -1036.1021 | 0.4289 | -1035.6722 | -0.0726 | -1035.7448 | -60.6 | / | / |
| Ser$^-$ | -1032.8983 | 0.3659 | -1032.5315 | -0.0705 | -1032.6020 | -57.2 | / | / |
| Ala$^-$ | -958.5191 | 0.3595 | -958.1586 | -0.0703 | -958.2288 | -59.7 | / | / |

[a] frequency calculations were performed at the level of structural optimization, HF-3c/CPCM

[b] gas phase energy levels were calculated from a separate set of optimized structures

[c] solvation energies of -6.32 kcal/mol and -103.45 kcal/mol for $H_2O$ and $H_3O^+$, respectively, were taken from literature[20]

[d] p$K_a$ values should be considered as "raw p$K_a$" and can be corrected by using linear correlations between calculated solvation free energies or p$K_a$ and known experimental values.[21] However, relative differences, in form of Δp$K_a$ values, can reflect the general trend sufficiently.
